# Co-translational assembly promotes functional diversification of paralogous proteins

**DOI:** 10.1101/2025.01.22.634331

**Authors:** Saurav Mallik, Angel F. Cisneros, Christian R. Landry, Emmanuel D. Levy

**Author notes:** To whom the correspondence should be addressed. S.M., C.R.L., E.D.L. Co-first authors (Saurav Mallik, Angel F. Cisneros). **Emails and ORCIDs** A.F.C.

## Abstract

Homomeric proteins are ubiquitous and mediate myriads of cellular functions. When a gene encoding a homomer duplicates, the resulting paralogs can either form distinct homomers, or evolve into a heteromer containing both paralogs. While such events have extensively shaped proteomes, the molecular mechanisms driving these fates and their associated functional consequences remain largely unknown. Here, we conducted a comprehensive phylogenomic analysis tracing gene duplication histories of 7,377 human paralogs across the eukaryotic lineage and identified their fates using protein interaction data. Simulations and data analyses show that cellular constraints must act as barriers to disfavor heteromerization and promote homomerization. We found that multiple cellular and molecular constraints can serve as barriers, including the lack of co-expression and co-localization. The main barrier, however, is co-translational assembly, which naturally promotes the self-assembly of each paralog from its corresponding mRNA, thus hindering heteromerization. We further established that heteromerization constrains functional divergence, with homomeric paralogs exhibiting stronger signatures of adaptive evolution and functional divergence compared to heteromeric paralogs. Together, these findings identify key biochemical and cellular properties that explain protein function diversification following gene duplication.

**One Sentence Summary:** Co-translational assembly drives the selective homo-oligomerization of paralogs, which in turn promotes their functional divergence.

**Graphical Abstract:** 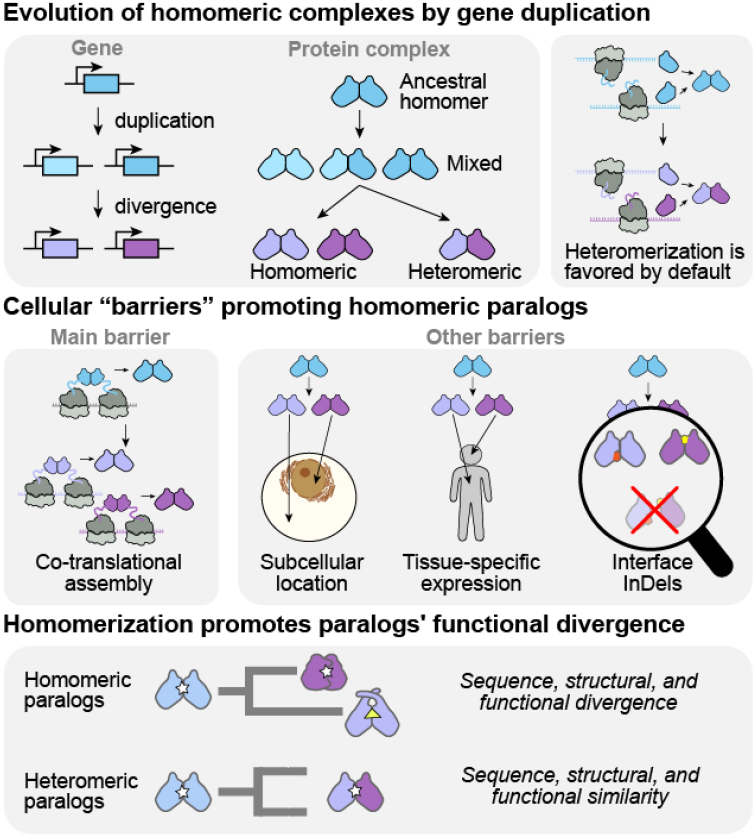

## INTRODUCTION

Gene duplication occurs frequently in eukaryotic genomes, arising through processes that range from the duplication of short chromosomal segments to that of entire chromosomes or genomes^1–5^. In principle, since the two duplicated copies (paralogs) are initially functionally redundant, one copy generally loses its function by accumulating mutations (*pseudogenization*) ^6–9^. Copies that survive may either retain their ancestral function (*conservation*) or diverge by splitting the ancestral function (*subfunctionalization*) or by acquiring new functions (*neofunctionalization*) ^10–17^.

Gene duplication is therefore a major source of functional innovation, which has prompted a large body of research to explore and rationalize the diverse evolutionary trajectories of paralogous pairs. These studies include the mathematical modeling of mutational and selection forces underlying paralog divergence ^4,10,17–20^, the experimental characterization of paralogs’ functional redundancies ^21–26^, or computational approaches characterizing their evolution and divergence in sequence and function ^27–31^. Although functional divergence appears to be key for the long term maintenance of paralogous pairs, many remain functionally redundant ^17,32,33^, which has important implications in cellular networks ^34^, development ^35–39^, immune system ^40^, neuronal regeneration ^41^, and cell cycle ^42,43^. For example, Tenascin is important for tissue remodeling and collagen fibrillogenesis; yet, its knockout does not lead to severe phenotypic defects in mice unless combined with knockouts of its paralogs ^37,41^. Similarly, knockout of protein tyrosine kinase *Src* does not block fibroblast cells’ G2 to mitosis transition, unless combined with the knockout of its paralogs *Fyn* and *Yes1* ^44^. Ultimately, the degree of functional divergence among paralogous pairs spans a wide spectrum, and general principles explaining why certain pairs diverge more than others remain elusive.

The evolution of a protein’s function can be constrained by multiple factors. For example, two proteins can become functionally coupled by direct physical interactions or via links in metabolic, regulatory, or signaling pathways ^45^. These couplings can restrict the evolutionary trajectories of paralogs, limiting the paths they would have otherwise taken if evolving independently ^45–49^. In particular, the dependency between subunits within a macromolecular complex can result in them being functional only in the oligomeric state. For example, this is seen in oligomeric enzymes in which the catalytic sites are shared between subunits ^50^, and also in large complexes such as the eukaryotic proteasome complex, where 14 paralogous subunits form the functional 20S core and are all essential for function _51_.

Selection for new functions during *neofunctionalization* and the loss of ancestral ones during *subfunctionalization* are key factors driving the maintenance of gene duplicates. Here, we asked whether structural and biophysical properties of proteins could influence these fates. We focused on the physical association between paralogs, which is systematically inherited from the duplication of homomer-encoding genes. Homomer-forming proteins are simple model systems for this study, because their duplications naturally introduce paralogs with or without direct cross-paralog interactions ^52,53^. In the absence of any constraints, paralogs originating from the gene duplication of a homomeric protein would form a mixture of homo- and heteromeric complexes ^49,54^ (*Mixed scenario*). Upon subsequent sequence divergence, this *Mixed* state might be retained. Alternatively, the two copies might lose their heteromeric interaction, resulting in two independent homomers (*Homomeric scenario*); or, they might lose the capacity to self-interact, resulting in a single heteromeric complex (*Heteromeric scenario*) ^55–59^. These alternative *fates* have been studied extensively, by experimental ^58–61^ or phylogenetic characterization of specific complexes ^62–65^, or by genomic analyses examining their relative abundances ^54–57^. Yet, the underlying molecular determinants of these *fates* and whether they associate with specific functional outcomes, remain largely unknown.

To address these questions, we conducted a comprehensive phylogenomic analysis tracing the gene duplication histories of human paralogous proteins across the eukaryotic lineage and examined their oligomeric fates using protein interaction data. This analysis along with simulations show that heteromerization is favored by neutral drift, unless constraints in the cellular environment act as barriers to the heteromeric interaction. We found that multiple cellular and molecular constraints can act as such barriers, including changes in gene expression profile, protein subcellular localization (spatiotemporal separation), or modifications at the oligomeric interfaces (structural incompatibility). Notably, co-translational assembly naturally induces each paralog to self-assemble from its respective mRNA and appeared as the barrier hindering paralog heteromerization the most. We also showed that direct physical interaction restricts the functional divergence of paralogous pairs. Such heteromeric paralogs tend to evolve in the nearly-neutral regime, accumulate substitutions and insertion/deletions more slowly than homomeric paralogs, and remain functionally more similar. By contrast, homomeric paralogs tend to evolve in the adaptive regime, accumulate evolutionary changes more rapidly, and increasingly diverge in structure and function with duplication age. Interestingly, the onset of their functional divergence appears to follow the apparition of the barrier to heteromerization. Together, these findings unveil a fundamental connection between co-translational assembly and paralog heteromerization, highlighting its widespread impact on the evolutionary origin of new protein functions.

## RESULTS

### Tracing the duplications of human paralogous genes across the eukaryotic tree of life

We traced gene duplication histories in the eukaryotic lineage leading to humans (*Methods*). To this end, we collected proteome sequences for 106 taxa representing all major eukaryotic clades; with an additional 11 *Proteobacteria* and six *Asgard archaea* proteomes that served as the outgroups (*Table S1*). We obtained the evolutionary relationships among these 123 organisms from the TimeTree database ^66^ and constructed an eukaryotic Tree of Life (which serves as the species tree, **Figure 1A**). We also adhered to the branch lengths obtained from TimeTree, which are species divergence times in million years, derived from molecular dating calibrated with geological records ^66^.

**Figure 1.**
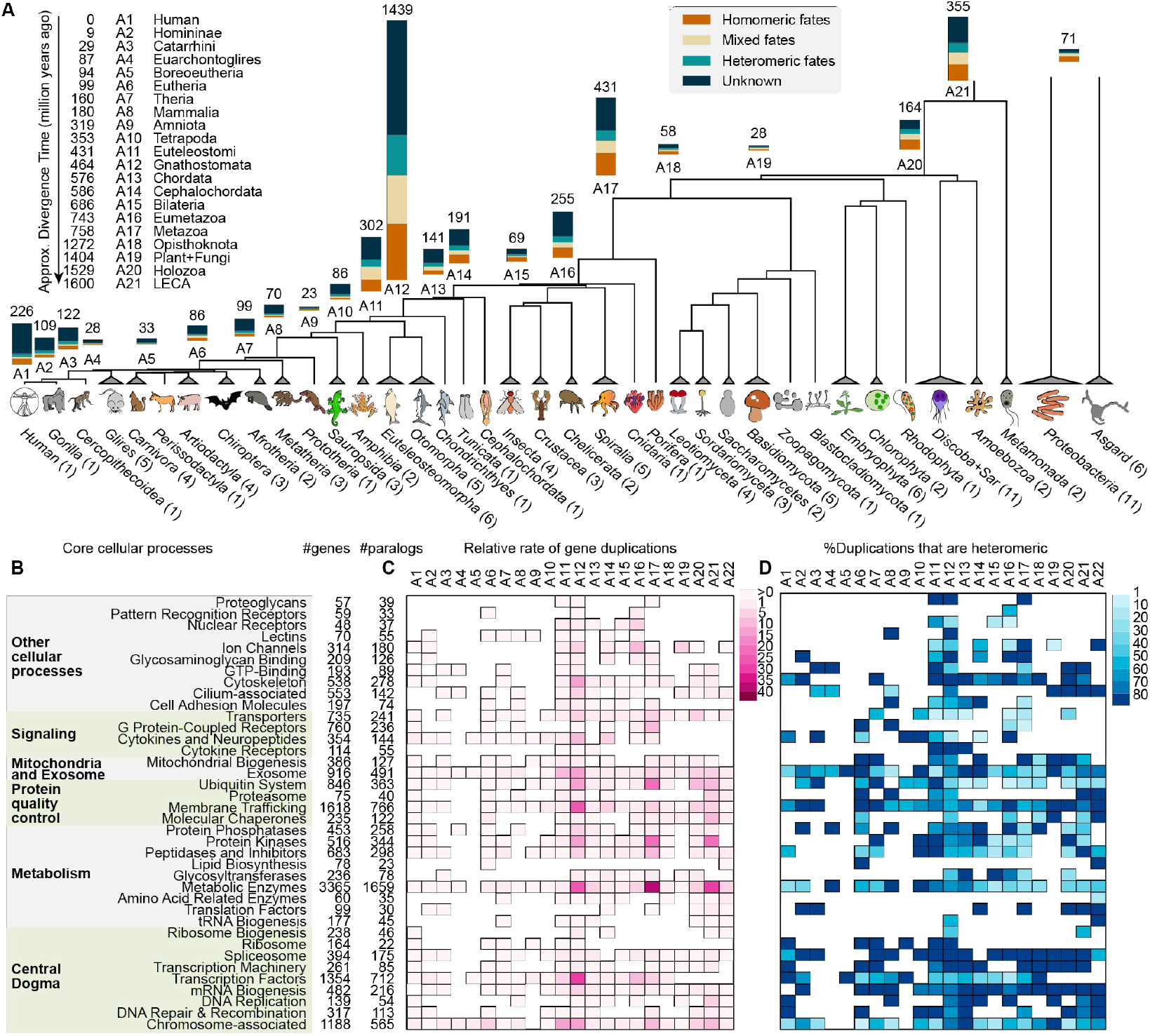
Inferring duplications of human paralogous genes across the eukaryotic tree of life. (**A**) The eukaryotic Tree of Life represents the phylogenetic relationships (obtained from the TimeTree database ^66^) among the 123 organisms (106 eukaryotes, 11 Proteobacteria, and 6 Asgard archaea) with fully sequenced genomes. Closely related organisms were grouped to represent the major phylogenetic clades and the #taxa are mentioned (*e*.*g*., *Xenopus laevis* and *Geotrypetes seraphini* were grouped into *Amphibia*). Leaves represent extant clades, while internal nodes (A1, A2, … A21) represent their common ancestors. Branch lengths are in *million years ago* (*mya*) unit, and reflect data from TimeTree. Organisms were sketched by the authors. The Last Eukaryotic Common Ancestor, or LECA (A21) represents the root of the eukaryotic sub-tree. Gene duplication events of human paralogs were phylogenetically inferred across the eukaryotic lineage. Vertical bars represent the frequency of inferred high- confidence duplication events at each ancestral node. Those giving rise to *Homomeric, Mixed*, and *Heteromeric* pairs in human cells are highlighted in orange, yellow, and green. Duplications for which the interaction status could not be reliably assigned are in dark blue. (**B**) Human protein-coding genes are classified into various functional categories according to KEGG annotations ^95^. For each functional category, the total number of annotated genes and the number of homomer-derived paralogous genes are shown (only those categories with ≥50 genes are plotted). (**C**) For each functional category, the relative proportion of gene duplications (normalized by the row-sum) across the ancestral nodes is shown as a heatmap. (**D**) Same as **C**, for the percentage of duplications that gave rise to a pair of heteromeric paralogs.

To detect human paralogous genes, we compared more than 2.1 million proteins (each corresponding to a gene) using all-*versus*-all BLAST ^67^ and then clustered them using a Markov cluster algorithm ^68,69^ into 60259 protein families. Out of those, 5330 families included ≥2 paralogous human genes.

To identify paralogous pairs that originated from homomeric ancestors, we examined their current oligomeric states. Molecular interaction data for human proteins were obtained from publicly available databases and include (i) crystallography, electron microscopy ^70^, and AlphaFold2-modeled structural data on protein complexes ^71^, (ii) subunit composition data for curated macromolecular complexes ^72,73^, and (iii) experimental protein-protein interaction data ^74–80^. These different datasets were integrated and filtered to minimize false positives, which yielded a high-quality set of 4,940 homomeric proteins and 142,226 heteromeric pairs (*Methods, Data S1*). Each paralogous pair was then classified into one of the three categories: *Homomeric, Heteromeric*, and *Mixed*. Paralogs featuring these oligomeric states are parsimoniously expected to have originated from ancestral homomers ^54–57^. We identified 1998 families including 7377 such paralogs (*Data S2*), highlighting that ∼35% (over a third) of the human proteome originates from ancestral homomers.

We used two complementary phylogenetic approaches to delineate the timing of gene duplication events (*Methods*). In the first approach, we inferred gene trees for each family and compared with the species tree to date the duplication events. In the second approach, given the gene copy numbers (GCN) for each species, we inferred duplication events for each family by a probabilistic reconstruction of the ancestral GCNs ^81^. Out of the 5381 inferred duplications, 4199 were considered high-confidence, as both approaches assigned the event at the same branch with ≥70 bootstrap support or ≥0.7 posterior probability (*Data S2*).

Our results showed that the human paralogous pairs duplicated at various stages of eukaryotic evolution (**Figure 1A**). About 33% of the duplications occurred in a single node *Gnathostomata* (A12, 1439 duplications), which coincides with the two/three rounds of whole-genome duplications during the origin of vertebrates ^1,82–85^. An additional ∼10% and ∼8% of the duplications occurred in *Metazoa* and LECA (A17 and A21, 431 and 355 duplications). Indeed, rampant gene duplications preceded the origin of eukaryotes ^86–91^ and metazoans ^92–94^, and the functional diversifications of the resulting paralogs are believed to have driven the complexification of eukaryotic cells and multicellular metazoan life ^86,92^.

What were the oligomeric *fates* of these duplicates? Since 1168 out of the 1998 families harbor nested duplications, the oligomeric fate of each duplication was inferred by parsimony (*Methods*). Results show that out of 4199 high-confidence duplications, 596, 640, and 983 gave rise to *Heteromeric, Mixed*, and *Homomeric* pairs respectively (**Figure 1A**, *Data S2*). The fate of 1980 duplications remained inconclusive due to the absence of experimental data on their physical interactions, and these were excluded from further analysis. Thus, ∼55% of the characterized high-confidence duplications resulted in the emergence of heteromeric paralogs (*Heteromeric*+*Mixed*), whereas the remaining ∼45% retained the ancestral homomeric state.

Next, we examined how these duplications contributed to the functional repertoire of mammalian proteomes. The human proteome was classified into different functional categories according to the KEGG database ^95^ annotations (**Figure 1B**, *Methods*). Homomer-derived paralogs contribute the least (∼21%) to functions related to translation machinery (transfer RNA biogenesis, ribosome biogenesis, and ribosome) and the most (∼70%) to lectins, nuclear receptors, proteoglycans, and kinases. Consistent with **Figure 1A**, three episodic bursts of duplications (*Gnathostomata, Metazoa*, and LECA) contributed to the expansion of most of these functional categories (**Figure 1C**).

We asked whether homo-versus heteromeric paralogs were associated with specific episodes of gene duplications. To that end, the fraction of duplications that gave rise to heteromeric pairs was computed for each branch of the species tree. However, no such coincidences were found and the fraction of heteromeric paralogs did not correlate with the frequency of duplication across the ancestral branches (Pearson R = 0.14, *Figure S1*). Overall, heteromeric fates dominate (>80%) in the following functional classes: DNA replication, repair, recombination, transcription, splicing, and proteasome, but are rare (<30%) in membrane proteins, chaperones, and glycosyltransferases (**Figure 1D**).

Taken together, these results highlight that over a third of the human proteome has descended from gene duplications of ancestral homomers. About 55% of the characterized duplication events gave rise to heteromeric paralogs, which are particularly frequent among functions associated with the central dogma.

### Barriers against paralog heteromerization drive the evolution of homomeric paralogs

What are the molecular determinants of different oligomeric fates? To address this question, we previously developed a biophysical model simulating the neutral accumulation of mutations in two duplicates and their impact on the probability that paralogs are maintained in homomeric *vs*. heteromeric states ^96^ (**Figure 2A**). These simulations were based on two important assumptions. First, that oligomerization occurs by random encounters of translated and folded monomers. Second, that mutations are fixed under a neutral model with respect to the three molecular complexes (AA, AB, and BB); specifically, (i) selection acts only on the total number of complexes without favoring the heteromer or either homomer, (ii) oligomerization interfaces remain conserved, and (iii) no new function emerges. For a dimeric complex, the first two assumptions lead to 50% heteromer and 25% of each homomer at binding equilibrium right after the duplication (**Figure 2A**). We found that neutral fixation of mutations to this *initial equilibrium* favors the enrichment of the heteromeric fates (*Heteromeric* or *Mixed*) at the end of the simulation (*final equilibrium*) for >95% of the cases examined ^96^. This is because due to their structural symmetry, the homomers accumulate destabilizing mutations (which are typically more abundant than stabilizing ones ^97,98^) faster than the heteromer. As this process continues, their binding affinities weaken, and larger fractions of the paralogs preferably heteromerize.

**Figure 2.**
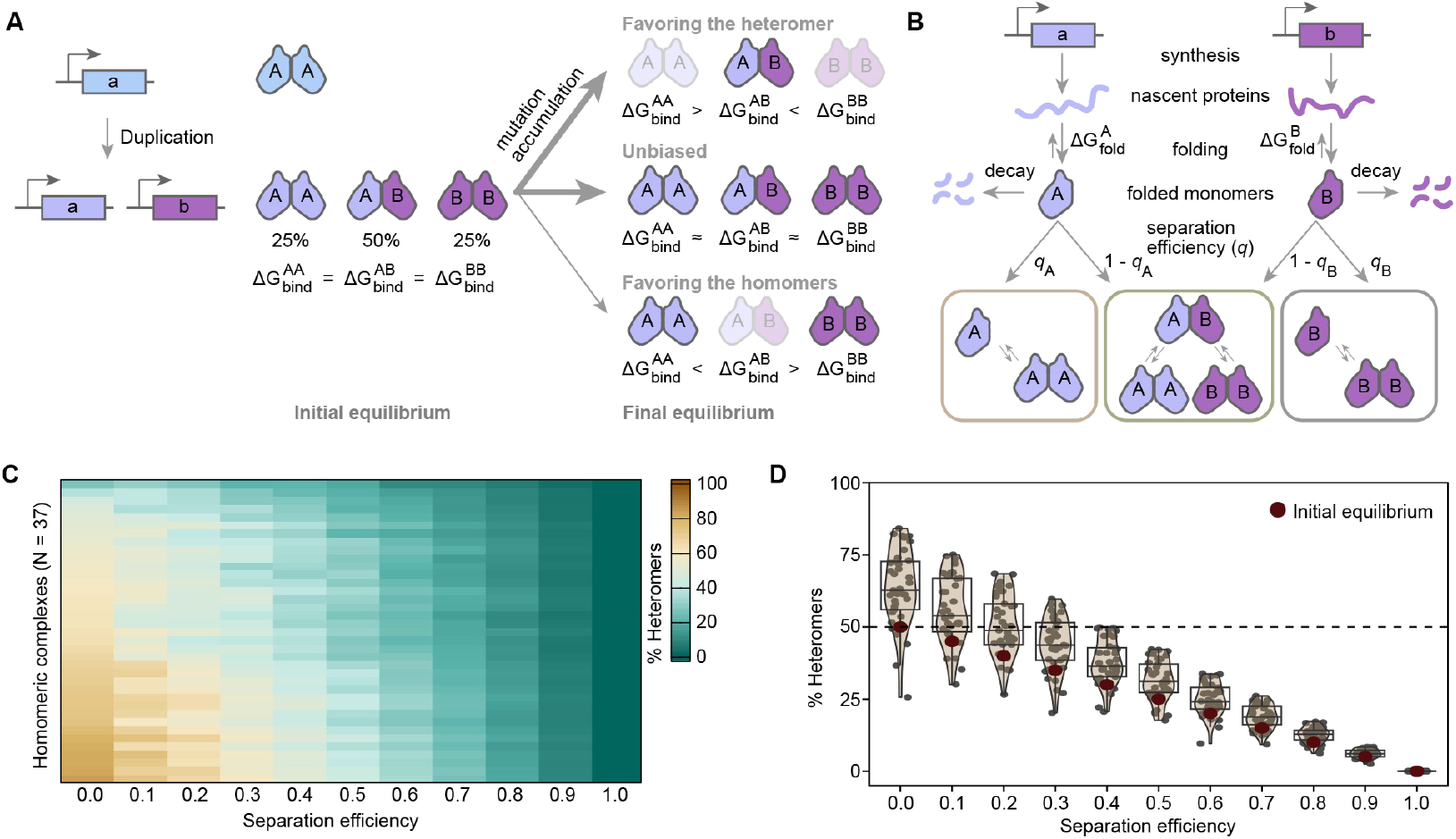
Barriers against paralog heteromerization drive the enrichment of *Homomeric paralogs*. (**A**) Duplication of a homomeric protein’s gene introduces a statistical mixture of homo- and heteromers. Assuming identical synthesis rates of each paralog, identical binding affinities of each complex, and stochastic oligomerization, the *initial equilibrium* includes 25% of each homomer and 50% heteromer. This changes upon further divergence, as the accumulating mutations alter these factors. Typically, mutational biases under a neutral model lead to the enrichment of the heteromer. (**B**) Simulation setup from Cisneros et al. ^96^, modified to include a barrier to paralog heteromerization. Each paralog is assigned a characteristic synthesis and decay rate and a folding free energy. The folded monomers are assigned a separation efficiency (*q*), which causes *q*% of them to only homomerize, while the remaining copies homo-/heteromerize by random encounters. The paralogs’ subsequent evolutionary divergence was simulated by introducing point mutations, with effects on folding free energies and binding affinities predicted by FoldX ^99,100^. These mutations alter the concentration of folded monomers and also the affinities of their homo/hetero-dimeric complexes. (**C**) This simulation framework shown in panel **B** was applied to 37 high-quality homodimers with crystallographic structures. The percentages of heterodimers obtained at the *final equilibrium* (median of 50 replicates, 200 mutations fixed for each) for different separation efficiencies are plotted as a heatmap. (**D**) The distribution of %heteromers in the *final equilibrium* for different separation efficiencies from panel C are plotted as violin plots. The %heteromers in the *initial equilibrium* are highlighted by red dots. The boxes and whiskers indicate the 25^th^-75^th^ percentiles and values that are 1.5 times the interquartile range.

Our observations are at odds with this model, since homomeric fates are observed with a higher proportion (∼45%, **Figure 1A**) than expected (<5%) from simulations, suggesting that one or many of the assumptions are not met in reality. These observations imply that underlying factors increase the likelihood of homomer formation. This could occur if certain barriers specifically hinder paralog heteromerization, causing homomers to become the dominant functionally active complexes in the *initial equilibrium*.

To reflect such barriers, we modified the simulation framework to add constraints (**Figure 2B**, *Methods*). Each paralog was assigned a *separation efficiency* (*q*) representing that *q*% of the folded monomers only formed homomers. The remaining folded monomers could form both homo- and heteromers by random encounters according to their respective binding affinities (Δ*G*_bind_). This *separation efficiency* thus acts as a barrier to heteromerization. As stronger *separation efficiencies* are applied, even if only on one paralog, heteromers are depleted in the *initial equilibrium* even if they have a stronger binding affinity than both homomers (*Figure S2*).

We applied this framework using various separation efficiencies ranging from 0 (no separation) to 1 (complete separation) to 37 homomeric structures (*Methods*) obtained from the Protein Data Bank (PDB) ^70^. These structures were sampled from major ECOD architecture types ^101^, and thereby represent independently evolved protein folds. In an iterative process, amino acid substitutions were introduced to the two paralogs that altered the Δ*G*_fold_ of their monomeric states and the Δ*G*_bind_ of their homo-/heteromeric complexes. Hence, the relative abundances of the mono- and oligomeric states changed upon each mutation. These mutations were randomly sampled from a pool of all possible single amino acid substitutions with effects on the folding free energies and the binding affinities estimated using FoldX ^99,100^. Each simulation was replicated 50 times, with each replicate continuing until 200 substitutions were fixed. For simplicity, identical separation efficiencies were assigned to each paralog.

Results show that as long as the barrier against paralog heteromerization was weak (0.0 ≤ *q* < 0.3), heteromers dominated the *final equilibrium* (**Figure 2C**). However, a moderate barrier (0.3 > *q* > 0.4) was sufficient to favor homomer evolution. For example, the homodimeric FMN-binding protein (PDB code: 3ZOF ^102^) forms 86% heteromers for *q* = 0.0, which reduces to 50% for *q* = 0.4 and to 30% for *q* = 0.7. Higher separation efficiencies further decreased the proportion of heteromers at the start of the simulation and reduced their maximum attainable concentration. When *q* = 1, the two paralogs could not interact at all, and no heteromer was formed. However, for *q* < 1, the two paralogs could form a heteromer from the beginning of the simulation. As simulation continued and mutations accumulated, the proportion of heteromer gradually increased relative to that in the beginning (*Table S2*), albeit these increases became smaller in magnitude with increasing values of *q* (**Figure 2D**).

In our simulations, the two paralogs may accumulate mutations at their oligomeric interfaces, but the overall interface patch remains at the same position on the structure. As such, these simulated paralogous pairs, even after evolving independently in the presence of a barrier, selection for the maintenance of homomerization makes them retain the ability to heteromerize if the barrier was lifted (*Table S3, Figure S3*). This is consistent with known paralogous pairs that localize to different subcellular compartments yet heteromerize *in vitro* ^74^.

Taken together, these results highlight that if oligomerization occurs by random encounters of monomers, then mutational biases combined with random genetic drift favors heteromerization of paralogs. Thus, the emergence of homomeric paralogs requires that barriers hinder the heteromeric interaction. With such barriers homomers can be favored, but what could these barriers be?

### Co-translational assembly is a major barrier promoting homomeric interactions

We systematically quantified the contribution of different protein properties that can act as barriers to heteromer formation, and thus promote homomeric paralogs in human cells. These barriers may have multiple independent origins and could either be acquired *de novo* or be inherited from the pre-duplication ancestor (**Figure 3A**). Acquired barriers may be physical, such as the two paralogs being located in different subcellular compartments, or being expressed in different sets of tissues and/or cell types ^56^. Alternatively, they could be temporal, as with a lack of gene co-expression ^56^. Acquired barriers can also be structural, such as InDels at oligomeric interfaces, which are known to efficiently modulate protein-protein interactions ^56,103,104^.

**Figure 3.**
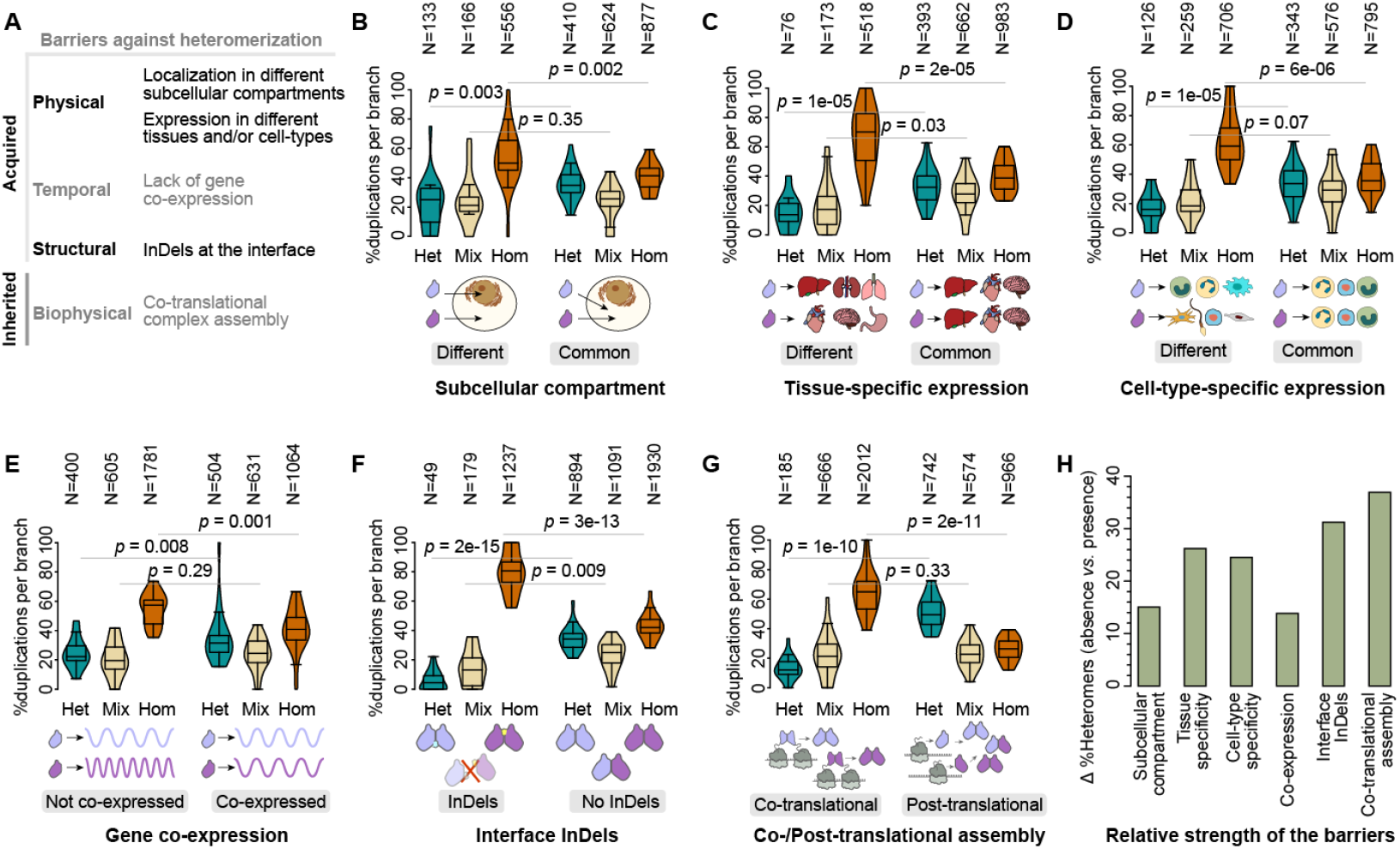
The effect of different barriers on the oligomeric state divergence of human paralogs. (**A**) Different types of barriers promoting homomeric paralogs. (**B-G**) Violin plots comparing the percentages of *Homomeric* (orange), *Heteromeric* (green), and *Mixed* (yellow) paralogous pairs, per family per ancestral branch, in the presence (left) *vs*. in the absence (right) of different barriers. The numbers represent the total number of corresponding duplication events across the eukaryotic lineage. Each violin includes 22 data points (each corresponds to an ancestral branch); violin features follow **Figure 2D**. Differences between the distributions were assessed by two- sample t-tests. The barriers are depicted as follows: (**B**) different *vs*. common (co-occurring in at least one) subcellular compartments, (**C**) expression in different *vs*. common sets of tissues given by intersection/union (*IoU*) in a total of 253 RNAseq-based tissue-specific expression datasets (*IoU* < 0.25 *vs. IoU* ≥ 0.25), (**D**) expression in different *vs*. common cell types (*IoU* < 0.25 *vs. IoU* ≥ 0.25, for a total of 1056 RNAseq based cell-type-specific expression datasets), (**E**) not coexpressed *vs*. coexpressed paralogs as measured by the Spearman correlation (*R*) of their mRNA abundances across 253 RNAseq-based tissue-specific expression datasets (*R* < 0.4 vs. *R* ≥ 0.4), (**F**) paralogs with *vs*. without interface InDels, (**G**) co-translational *vs*. post-translational assembly. (**H**) Barplots compare the strengths of different barriers as the deviation (Δ) of the average %*Homomeric* paralogs (per family per branch) observed in its absence *vs*. its presence.

To quantify the effects of acquired barriers, we integrated multiple types of omics data together with the reconstructed histories of the duplicates (*Methods*). Protein subcellular localization as well as tissue and cell-type-specific gene expression data (*Data S3*) were obtained from the Human Protein Atlas ^105^. We first examined paralogous pairs that co-localize in the same common subcellular compartment *vs*. those that do not. For each category, the percentages of *Homomeric, Heteromeric*, and *Mixed* pairs per family were computed for each ancestral branch of the species tree. We found that *Homomeric* and *Heteromeric fates* show opposite trends. *Homomer* representation decreases from 54% among paralogous pairs that do not co-localize in the same subcellular compartment to 41% among pairs that do co-localize (*p* = 0.002, **Figure 3B**). By contrast, the *Heterome*r representation correspondingly increases from 22% to 37% (*p* = 0.003, **Figure 3B**).

Similar trends were observed for paralogous pairs expressed in different *vs*. common sets of tissues (*Homomer* representation decrease from 66% to 40%, *p* = 2e-05, **Figure 3C**) and cell-types (*Homomer* representation decrease from 62% to 38%, *p* = 1e-05, **Figure 3D**). Also for temporal barriers we found that *Homomer* representation decreases from 55% among weakly co-expressed paralogous pairs to 41% among strongly co-expressed pairs (*p* = 0.001, **Figure 3E**).

To examine the role of structural barriers, we used AlphaFold Multimer ^106,107^ to model quaternary structures of homo- and heteromeric paralogs (*Methods*). A total of 3760 homomeric and 955 heteromeric model structures (*Data S4*) were examined to identify paralog-specific InDels at the oligomeric interfaces (both small InDels and domain gain/losses). We found that *Homomer* representation decreases from 76% among paralogs with interface InDels to 44% among those without (*p* = 3e-13, **Figure 3F**). In the same categories, the *Heteromer* representation correspondingly increases from 6% to 34% (*p* = 2e-15, **Figure 3F**).

Importantly, alterations of gene expression or protein subcellular localization are not expected to be effective immediately after the duplication. Therefore, the pronounced increase in homomer representation in the presence of these barriers could be the consequence of the fate rather than the driver of the fate (oligomeric state divergence first *versus* barrier first scenarios).

However, we reasoned that another barrier, co-translational assembly, would exist immediately after duplication. Co-translationally assembling homomers begin to interact as they emerge from their respective ribosomes ^108–110^. For homomers, when both ribosomes translate the same mRNA (*cis*-assembly), this mechanism creates a natural barrier to heteromerization. In this scenario, two paralogs are synthesized as two independent homomers even if they are co-expressed and co-localized in the same subcellular compartment. Moreover, we recently showed that co-translational assembly originates in structural and biophysical features of protein complexes and typically involves subunits that are geometrically intertwined and partner-stabilized ^111^. Because these subunits tend to be unstable in isolation, we expect the paralogs’ *post facto* heteromerization by subunit exchange to be rare as well.

Bertolini et al. ^112^ identified thousands of human proteins undergoing co-translational assembly and these could be accurately predicted from their structural characteristics ^111^. Based on this integrated dataset, ∼17% of the human proteome was found to assemble co-translationally and ∼53% was annotated as assembling post-translationally. Projecting the reconstructed history of duplicates onto these data revealed that *Homomer* representation decreased from 67% among co-translational paralogs to 28% among post-translational paralogs (*p* = 2e-11, **Figure 3G**). In the same categories, the *Heteromer* representation increased from 13% to 49% (*p* = 1e-10, **Figure 3G**). This result provides conclusive evidence that co-translational self-assembly is a major determinant of paralogs’ homomeric fates.

To compare the strength of barriers we used the difference (Δ) between the average %*Homomeric fates* in the presence *vs*. absence of a barrier (**Figure 3H**). We found that the temporal barrier (lack of gene co-expression) was the weakest, likely because human cells include ∼1000-fold variation of protein half-lives ^113–115^ and so protein products of weakly co-expressed genes can still interact if they happen to be long-lived. Physical barriers are of intermediate strength (tissue/cell-type-specific expression being stronger than subcellular localization). However, co-translational assembly appeared to be the strongest while interface InDels (structural barrier) was the second strongest.

Taken together, these results provide conclusive evidence that homomeric paralogs dominate when certain barriers specifically hinder paralog heteromerization. In particular, co-translational assembly, originating from the structural features of protein complexes, appeared as the strongest barrier promoting homomeric paralogs in human cells and throughout eukaryotic evolution.

### Heteromerization constrains paralogs’ sequence, structural, and functional divergence

To examine whether heteromerization of paralogs constrains their functional divergence, we compared homo- and heteromeric paralogs using three complementary measures of functional divergence (*Methods*). First, for paralogous enzymes we analyzed their Enzyme Commission (EC) numbers, which represents the biochemical reaction they catalyze. EC number differences thus represent altered substrate-specificity or reaction chemistry ^116^. EC numbers of 3758 human enzymes obtained from Expasy ^116^ and KEGG ^95^ were mapped to 1622 paralogous pairs (*Data S3*). We found that the average frequency of unique EC numbers per family (*i*.*e*. one paralog catalyzes a reaction the other one doesn’t) increases with duplication age more rapidly for *Homomeric pairs* than for *Heteromeric* and *Mixed* (t-test *p* < 2E-05, **Figure 4A**).

**Figure 4.**
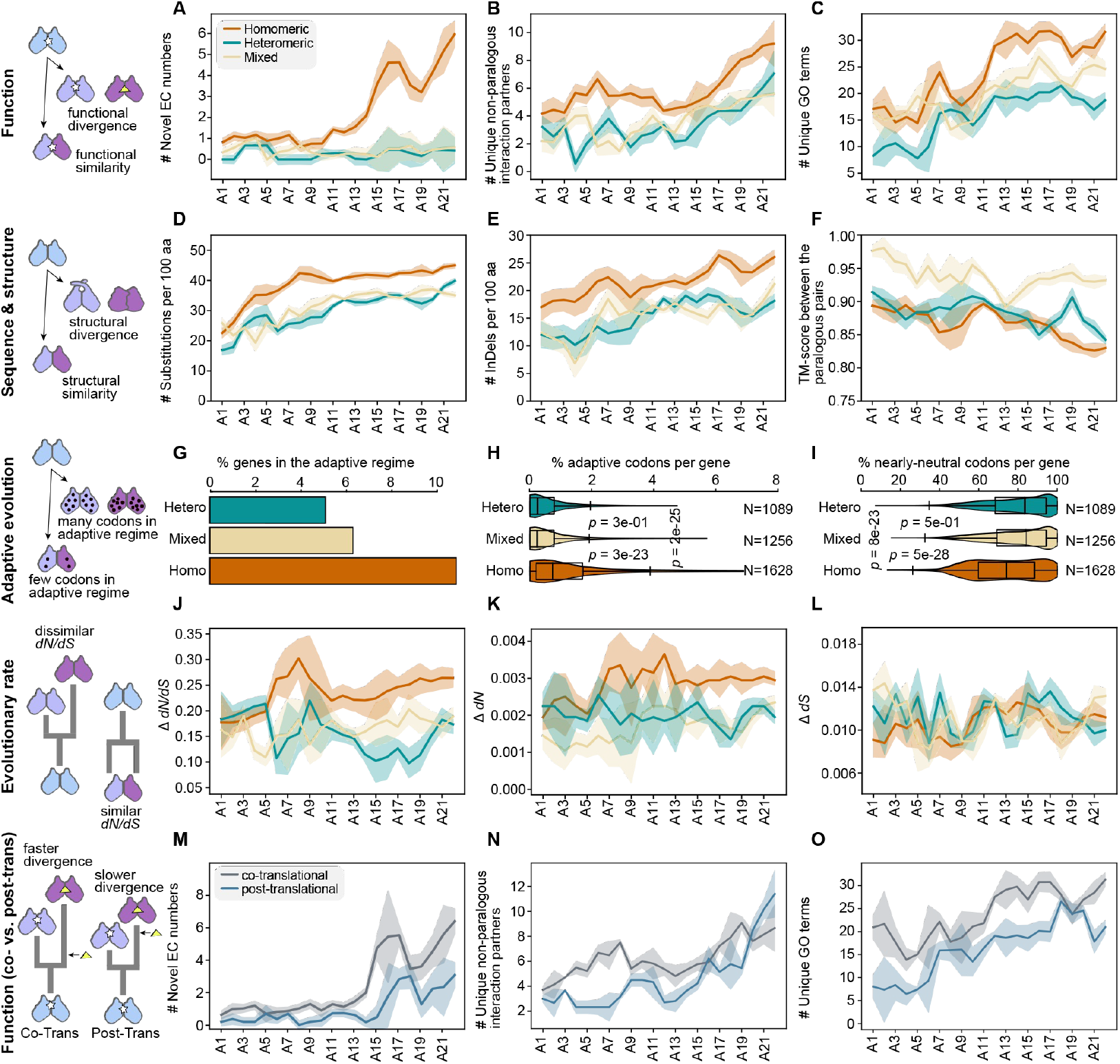
Homomeric paralogs increasingly diverge in sequence, structure, and function with duplication age. (**A**-**C**) Ribbon plots represent the functional divergence of *Homomeric* (orange), *Heteromeric* (green) and *Mixed* (yellow) paralogous pairs. The lines and ribbons represent the mean and standard deviations. For clarity, each curve was smoothed using a sliding window approach (window size = 3). (**A**) The count of novel EC numbers per family. (**B**) The count of novel non-paralogous PPI partners per family. (**C**) The count of novel GO categories per family. (**D-F**) Same as panels **A-C**, for sequence and structural divergence of paralogous pairs. (**D**) The average frequency of substitutions, per 100 amino acids, per family. (**E**) The average frequency of InDel sites, per 100 amino acids, per family. (**F**) The average TM-score (of the structural superposition) between AlphaFold2-modeled structures of paralogous pairs, per family. (**G**-**I**) Genes/codons are assigned to evolve in adaptive or nearly-neutral regimes by comparing their nonsynonymous (*dN*) *vs*. synonymous substitution rates (*dS*). (**G**) Bar plots depicting the %paralogous genes in the adaptive regime. (**H**) Violin plots comparing the %codons per gene in the adaptive regime. Differences between the distributions were assessed by two-sample t-tests. (**I**) Same as panel-**H**, for the %codons per gene in the nearly-neutral regime. (**J**-**L**) Same as panels **A-C**, for the difference (Δ) of evolutionary rates (*dN*/*dS, dN*, and *dS* are plotted along the Y-axis in panels **D, E** and **F**). (**M**-**O**) Same as panels **A-C**, for functional divergence of co- (gray) and post-translational paralogs (blue).

Next, we examined how paralogous pairs diverge to interact with unique unrelated partners. To that end, we mined 22184 heteromeric PPIs between co-complex subunits ^72,73^ (*Data S1*), which included interaction data for 1480 paralogous pairs. The average frequency of unique interaction partners per family (*i*.*e*. one paralog interacts with an unrelated partner the other one doesn’t) also increases with duplication age more rapidly for *Homomeric pairs* than *Heteromeric* and *Mixed* (*p* < 8E-07, **Figure 4B**).

Finally, Gene Ontology (GO) data was analyzed to explore how paralogous pairs diverge to perform new functions. Here, GO terms related to ‘biological process’ and ‘molecular function’ ^117,118^ were obtained for 19070 human proteins and mapped onto 2245 paralogous pairs (*Data S3*). The average count of novel GO terms per family (*i*.*e*. one paralog is assigned a GO term that the other one isn’t) also increases with duplication age more rapidly for *Homomeric* pairs than *Heteromeric* and *Mixed* (*p* < 3E-04, **Figure 4C**).

We asked whether the accelerated functional divergence of homomeric paralogs is coupled with their rapid sequence and structural divergence. Analyzing pairwise sequence alignments of paralogous pairs (*Methods*) we found that accumulation of substitutions and InDels increases with duplication age more rapidly for *Homomeric pairs* than for *Heteromeric* and *Mixed* (*p* < 3E-06, and *p* < 2E-08, **Figure 4D-E**). Furthermore, homomeric paralogs accumulate InDels and substitutions both at the interface and surface, whereas heteromeric paralogs predominantly keep their interfaces conserved (*Figure S4*). This accelerated InDel accumulation is consistent with a faster divergence of protein structure and function in homomeric paralogous pairs ^56,103,104,119–122^. Indeed, assessing structural divergence of paralogous pairs by the template-modeling score (TM-score ^123^) shows that *Homomeric* pairs diverged more rapidly than *Heteromeric* and *Mixed* pairs (*p* < 0.01, **Figure 4F**). Interestingly, *Mixed* pairs remained structurally more similar than *Heteromeric* (*p* < 3E-08). Together, these results highlight that the rapid functional divergence of homomeric paralogs is associated with their accelerated sequence and structural divergence.

Rapid accumulation of substitutions and acquisition of new functions are often driven by positive selection. Hence, we compared the nonsynonymous (*dN*) and synonymous substitution rates (*dS*) of paralogous genes seeking signatures of positive selection or neutral drift. The ratio of the two, called *ω* (or *dN*/*dS*), provides a measure of the strength of selection exerted at the amino acid level ^124,125^. In principle, *ω* is expected to be =1, <1, or >1 under neutral, purifying, or positive selection scenarios ^126–128^. However, in practice, most proteins evolve under a mix of positive and purifying selection and are dominated by the latter, typically leading to *ω* < 1 even in the presence of positive selection. A more sensitive approach is to compare the observed *ω* against that expected under an explicitly neutral model (*ω*_*0*_). Indeed, *ω > ω*_*0*_ and *ω* ≈ *ω*_*0*_ highlight evolution in the adaptive and nearly-neutral regimes.

We used a dataset ^129^ annotating the evolution regime (*e*.*g*., adaptive and nearly-neutral) of genes and codons in the human genome. At the gene level, significantly more homomeric paralogs (∼10% *versus* ∼6% for heteromers) were observed to evolve in the adaptive regime (**Figure 4G**). Extending our analysis to the codon level, we found that *Homomeric* paralogs showed a higher percentage of codons in the adaptive regime when compared to the *Heteromeric* and *Mixed* categories (*p* < 3e-25, **Figure 4H**). Conversely, *Heteromeric* and *Mixed* paralogs harbored a higher percentage of codons in the nearly-neutral regime when compared to the *Homomeric* category (*p* < 8e-23, **Figure 4I**). These results highlight that homomeric and heteromeric paralogs are enriched in genes and codons evolving in the adaptive and nearly-neutral regimes respectively.

We asked whether the functional divergence of *Homomeric* pairs is coupled to a pronounced sequence divergence. To that end, we compared the difference (Δ) of *ω* for homo- and heteromeric paralogs. Results show that *Homomeric* pairs typically display significantly higher differences (*i*.*e*. more dissimilar *ω*) than *Heteromeric* and *Mixed* pairs of the same duplication age (*p* < 2E-10, **Figure 4J**). These trends were recapitulated for *dN* (*p* < 4E-08, **Figure 4K**) but not for *dS* (*p* > 0.2, **Figure 4L**), highlighting that they correspond to changes in the protein sequence, as previously reflected in **Figure 4D-E**. Together, these results highlight that homomeric paralogs tend to evolve more asymmetrically (functional divergence plus dissimilar evolutionary rates) than heteromeric paralogs.

Does the asymmetric evolution of *Homomeric* paralogs kick in immediately after the duplication? As depicted in **Figure 4A-C** and **J-L**, it appears that pronounced differences in EC numbers, interaction partners, molecular functions, and evolutionary rates are primarily found between homo- and heteromeric paralogs that duplicated before the *Boreoeutheria* node (node A5, placental mammals, originated 94(±5) million years ago in mid-Cretaceous ^66,130^). Hence, a ‘lag phase’ of ∼100 million years appears to exist between the duplication event and the onset of asymmetric evolution. Interestingly, as depicted in **Figure 4M-O**, this lag is relatively shorter in co-translational homomeric paralogs (inherited barrier, effective immediately) as compared to post-translational (acquired barriers, require evolutionary change accumulation). Thus, the onset of asymmetric evolution appears to follow the apparition of the barrier to heteromerization.

These results show that the asymmetric evolution of paralogous pairs is tied to their structural and biophysical independence. Homomeric paralogs diverge more rapidly over time in enzymatic reactions, interaction partners, and molecular functions, while heteromeric paralogs remain more functionally linked.

## DISCUSSION

We have shown that gene duplication of homomeric proteins often results in independent homomers if heteromerization is prevented by barriers. Co-translational self-assembly of each paralog appeared as an important such barrier. Interestingly, this barrier likely operates immediately after duplication because the structural properties that drive co-translational assembly are inherited from the ancestor. By contrast, other barriers such as spatiotemporal separation or structural incompatibility likely appear with a delay after the duplication event. In the absence of such barriers, the heteromer is favored. As they accumulate mutations, both paralogs could become ‘entrenched’ into a heteromeric complex, which would slow down their functional divergence ^61,131–133^. This phenomenon, known as ‘constructive neutral evolution’ ^134^ is likely widespread in eukaryotes ^135,136^ and could have contributed to homomer-dominant prokaryotic proteomes transforming into heteromer-dominant eukaryotic proteomes ^55,56^.

Co-translational self-assembly appeared as the most significant barrier promoting homomeric fates (**Figure 3G,H**). In principle, co-translational self-assembly can occur either in *cis* or in *trans*; that is, from two ribosomes translating the same or different mRNAs respectively. While a handful of experimentally characterized homomers were shown to assemble in *cis* ^112,137,138^, which mode of assembly dominates in the cell, *cis* or *trans*, remained elusive. Our results are consistent with the dominance of *cis*-assembly in eukaryotic genomes, and this trend was also reflected in co-translational homomeric proteins being depleted in mutations associated with autosomal dominant disorders ^137^.

Many paralogs are known to evolve asymmetrically (*i*.*e*. diverge in structure, function, and evolutionary rates), but what fraction of paralogs do so and why, is debated ^139–144^. We showed that asymmetric evolution is coupled with paralogs’ structural and biophysical independence. These results are complementary to observations made for yeast whole-genome duplicates ^26^.

In summary, we unveiled a fundamental connection between co-translational assembly and the evolutionary divergence of protein oligomeric state and function. The resulting framework highlights the power of combining phylogenomics with multi-OMICs data and further helps resolve long-standing questions such as why paralogs appear to reach specific oligomeric fates, why only some paralogs evolve asymmetrically, and why temporal lags exist between whole genome duplications and subsequent adaptive radiation.

## Acknowledgments

We thank Yitzhak Pilpel, Sudip Kundu, Soham Dibyachintan, and Carla Bautista Rodríguez for helpful discussions. E.D.L. acknowledges support from the European Research Council under the European Union’s Horizon 2020 research and innovation program (grant agreement No. 819318), the Human Frontier Science Program (ref. no: RGP0016/2022), the Israel Science Foundation (grant No. 1452/18), and the Abisch-Frenkel Foundation. C.R.L. acknowledges support from Canadian Institutes of Health Research (CIHR) Foundation grant No. 387697, NSERC discovery grant, Canada Research Chair in Cellular Systems and Synthetic Biology. A.F.C. acknowledges support from the Fonds de recherche du Québec - Nature et technologies (dossier 290237), the Ministère de l’Enseignement Supérieur du Québec, the Agencia mexicana para la cooperación y el desarrollo internacional, a Mitacs Globalink Research Award (award number IT28316), and a PROTEO Graduate Scholarship.

## Author contributions

S.M., A.F.C., C.R.L., and E.D.L. conceived and conceptualized the research. S.M. performed the phylogenetic, OMICs, and evolutionary rate analyses shown in Figures 1, 3, and 4. A.F.C. performed the biophysical simulations shown in Figure 2, structural analysis shown in Figure 3G and 4F, and conceptualized the barrier strength analysis shown in Figure 3H. A.F.C. also generated the AlphaFold2 models of heteromeric paralogs. S.M. and A.F.C. prepared the figures and wrote the first draft. E.D.L. and C.R.L. acquired funding and supervised the study. All authors discussed and commented on the results and edited versions of the paper.

## Data and Materials availability

All data used or generated in this study are provided as Supplementary Materials. AlphaFold2 models of paralogous protein complexes are available in Zenodo: https://doi.org/10.5281/zenodo.14158203.

## Materials and Methods

### Taxon sampling and species tree construction

We sampled 123 organisms with fully sequenced and annotated genomes (*Table S1*) following our previously published methodology ^145^. Briefly, for all major phylogenetic clades of the domain Eukaryota, one or more representative species were selected, which included a total of 106 taxa. Aiming to uniformly sample the Eukaryota, we ensured that any pair of species in our dataset has a minimum divergence time of 65 million years, with divergence time data obtained from TimeTree v5 ^66^. In addition, we recruited 17 outgroup species, comprising 11 and 6 species representing various *Proteobacterial* and *Asgard archaea* classes respectively.

The TimeTree database v5 comprises the phylogenetic relationships among 137306 species and their divergence times estimated by the molecular clock analyses calibrated by fossil records ^66^. Our species set was submitted to the TimeTree database to obtain the eukaryotic Tree of Life which served as the species tree. The tree leaves represent the extant species and the internal nodes represent their now-extinct ancestors. The tree topology was manually adjusted to include the *Proteobacterial* and the *Asgard archaea* clades as the outgroups since their endosymbiosis gave rise to the first eukaryote ^146^. The branch lengths represent the evolutionary divergence times (million years) as documented in TimeTree. Due to the near-absence of fossil records, for ancestral branches older than *Metazoa*, the divergence time estimates are based solely on molecular dating and are therefore putative.

### Genome-wide comparative phylogenomic analysis

#### Identifying protein families

Annotated proteome sequences of the 123 organisms were obtained from UniProt ^147^, including each gene’s canonical protein sequence. Gene families were identified using the OrthoFinder package v2.5.5 ^69^. OrthoFinder analyses include two steps. In the first step, protein sequences were compared in all-*vs*.-all BLASTp searches ^67^. The resulting BLASTp hits were filtered by removing all query-subject-pairs for which the alignment coverage was <30% of either the total query or subject sequence length. This step is expected to significantly improve phylogenetic trees and orthology inference in the subsequent steps ^148,149^. In the second step, based on these modified BLASTp tables the remaining 2162331 sequences were clustered (inflation parameter of 1.5) into 60259 protein families which were considered proxies of protein families.

#### Filtering protein families that have descended from homomeric ancestors

A total of 5330 families, out of the initial 60259, included at least two human paralogs. To identify which of these families are descendants of ancestral homomeric proteins, we examined the molecular interaction status of the human paralogs. To that end, a curated dataset of 4940 homomeric and 142226 heteromeric interactions (see “**Compiling molecular interaction data**”) was mapped to these paralogous pairs. Each paralogous pair was classified into one of the three categories, *Homomeric* (both paralogs are homomeric and they do not heteromerize), *Heteromeric* (the two paralogs form a heteromer, and no homomeric interaction was observed), or *Mixed* (both paralogs are homomeric and they also form a heteromer). Paralogous pairs of these three categories are parsimoniously expected to have emerged from ancestral homomeric proteins ^54–57^. We identified 1998 protein families, each including at least one such paralogous pair (*Data S2*). These 1998 families include 7377 human proteins.

#### Multiple sequence alignment and trimming

Protein sequences of each family were aligned by the 3D-Coffee alignment tool, implemented in T-Coffee package ^150^. 3D-Coffee generates structure-guided sequence alignments. Briefly, 3D structures of multiple template proteins are aligned to create a multiple structural alignment, which is then used as a guide to align the remaining sequences. We collected proteome-scale AlphaFold2-modeled ^151^ monomeric structures of 10 organisms (*Homo sapiens, Mus musculus, Danio rerio, Drosophila melanogaster, Caenorhabditis elegans, Arabidopsis thaliana, Saccharomyces cerevisiae, Plasmodium falciparum, Trypanosoma brucei*, and *Escherichia coli*). These structures were used as the templates. These multiple alignments were subsequently trimmed using the trimAL tool ^152^ (parameter settings: -gt 0.6 -cons 50) to remove gap-majority columns.

#### Phylogenetic tree inference

For each of the 1998 protein families, command-line Mega v11.0.8 was used to infer Maximum-Likelihood unrooted protein trees (model = LG+F, Rates among Sites = G+I, number of discrete gamma categories = 5, ML Heuristic Method = SPR level 5, Initial Tree = NJ/BioNJ, Test of Phylogeny = 100 Bootstrap Replicates). These trees are available in newick format in *Data S2*.

### Compiling molecular interaction data

#### Complexes with experimental/predicted structures

We obtained structures of 837 human homomeric complexes resolved by X-ray crystallography or Electron Microscopy (EM) from the 3DComplex database ^153,154^, based on the following criteria. (i) We selected homomers whose QSBio error rate was below 15% ^153^. (ii) We imposed a minimum sequence coverage of ≥ 70% for the respective UniProt sequence ^147^. (iii) Structures corresponding to artificial fusion proteins (N = 33) or proteins encoded by transposon elements (N = 12) were discarded.

Additionally, we used the results from our recently developed AlphaFold2-based ^106^ framework that predicts homomeric structures across full proteomes ^71^. These predictions enabled us to map a high-quality dataset of homomeric complex structures for 3584 human proteins.

For heteromers, we collected a set of 2606 heteromeric complex structures from Protein Data Bank ^70^, from which subunit composition and inter-subunit interaction data were extracted.

#### Curated macromolecular complexes

Subunit protein composition data for 1265 and 2303 experimentally annotated human heteromeric complexes were obtained from Complex Portal ^72^ and hu.MAP v2.0 ^73^ databases.

#### Protein-protein interaction data

Human protein-protein interaction (PPI) data, including both homo- and heteromeric interactions were obtained from seven databases. This initial dataset included 73834, 142706, 64923, 181555, 51503, 114654, and 173350 interacting pairs obtained from BioGrid ^74^, IntAct ^75^, HuRI ^76^, Mentha ^77^, MINT ^78^, HiNT ^79^, and HIPPIE ^80^. In total, these databases included 390709 unique PPI interactions. Since PPI data tend to be relatively noisy, we subsequently filtered these interactions as described previously in ref. ^55^. Briefly, in the first step, all predicted interactions and text-mining-based interactions were removed. In the second step, to minimize false positives, we demanded that the interaction between two proteins be observed using both proteins as bait and as prey. These filtering steps provided a high-quality dataset of 192543 interactions.

#### A comprehensive human interactome

We aimed to unify these different datasets together to obtain a comprehensive set of human homomeric proteins and heteromeric protein pairs. A homomeric interaction was included in the final dataset (i) if the interaction was observed in at least one X-ray crystallographic or EM or AlphaFold2-predicted structure, or (ii) if the interaction was observed in at least three independent PPI experiments. These filtering steps provided a dataset of 4940 human homomeric proteins (*Data S1*).

A heteromeric pair was included in the final dataset, (i) if their pairwise interaction was observed in at least one X-ray crystallographic or EM or AlphaFold2-predicted structure, or (ii) if both are members of the same curated complex, and their pairwise interaction was observed in at least one PPI experiment, or (iii) if their pairwise interaction was observed in at least three independent PPI experiments. These filtering steps provided a final dataset of 142226 human heteromeric protein pairs (*Data S1*).

### Annotating gene duplication events

We annotated gene duplication events on the species tree by two complementary approaches.

#### Gene tree-species tree reconciliation approach

We used Notung v2.9 ^155^ to reconcile the protein trees of the 1998 protein families with the species treeusing the duplication-transfer-loss model (-- rearrange --threshold 70.0 --costdup 1.5 --costtrans 1.0 --costloss 1.0 --prune --treeoutput nhx -- events). Given a protein tree and a species tree, Notung infers gene duplications, losses, and transfers along a species tree. For each inferred duplication event, the bootstrap support of the respective branch in the protein tree was considered as the statistical confidence of the inference.

#### Maximum-Likelihood-based inference of ancestral gene copy numbers

We annotated gene duplication events using a complementary approach which is based on inferring the gene copy numbers for the ancestral nodes of the species tree. We used PastML ^81^, which infers ancestral characters on the internal nodes of a rooted phylogenetic tree (species tree) with annotated tips (copy-number annotation) using maximum-likelihood methods (ML method = Marginal Posterior Probabilities Approximation or MPPA, character evolution model = Estimate-from-Tips or EFT). A posterior probability score is assigned to each copy-number inference, which was considered as the statistical confidence for inferring duplication events.

For the 1998 families, we inferred a total of 5381 duplication events, which were subsequently filtered based on the following criteria. First, we only considered those duplications for which Notung-based and PastML-based inferences converged, *i*.*e*., they both traced the duplication event to the same ancestral branch. Additionally, only high-confidence duplication inferences (those inferred with ≥ 70 bootstrap support or with ≥ 0.7 posterior probability) were kept. These filtering steps provided a final set of 4386 high-confidence duplications (*Data S2*).

#### Assigning the oligomeric fates of ancestral duplications

Given the oligomeric states of the extant human paralogous genes, the fates of ancestral duplications were assigned by maximum parsimony using PastML ^81^. The results are available in *Data S2*.

### Simulating Oligomeric State Evolution

#### Selection of protein structures

A total of 37 structures obtained from Protein Data Bank (PDB) ^70^ was analyzed (*Table S2, S3*); this dataset was compiled in our previous work ^96^. Briefly, we mined PDB for homodimeric structures resolved by X-ray crystallography composed of two identical subunits 60-450 amino acids in length. Structures were sampled from major ECOD architecture types ^101^, each of which represents an independently evolved protein fold. To remove redundancy, we clustered the resulting 104 structures using CD-Hit version 4.8.1 ^156,157^ at 40% sequence identity. All structures were identified in different clusters. Finally, we selected 37 structures that were annotated in the highest quality tier in the QSBio database ^153^ (error probability < 2%).

#### Estimation of mutational effects

Effects of point mutations on each homomeric structure were estimated as in our previous work ^96^. Briefly, the biological assemblies of the 37 structures were energy-minimized using the FoldX Repair function ^99^ ten times to ensure convergence to an energy minimum ^158^. Then, *in silico* mutagenesis was performed using the FoldX BuildModel and AnalyseComplex functions ^99^ with the MutateX workflow ^100^. The effects of mutations on the folding free energy and the binding affinity of heterodimers were calculated by simulating mutations on only one of the protein copies. Effects on the binding affinity of the homodimer were obtained by simulating mutations on the two copies.

#### Framework for simulations

Simulations were initialized with a gene encoding a homomeric protein with typical values of folding free energy ^159,160^ and binding affinity ^161,162^. Folding free energy is used to calculate the fraction of properly folded subunits following equation 1 ^163^:

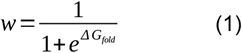

where *w* is the fraction of folded proteins and Δ*G*_*fold*_ is the free energy of the native fold.

Properly folded subunits were then assigned a separation efficiency (*q*), representing the fraction of protein copies that can only form homomers. The rest of the protein copies were allowed to mix, thus forming both homo- and heterodimers based on their binding affinities. Equilibrium concentrations of monomers, homodimers, and heterodimers were then calculated using the same system of equations as in our previous work ^96^. Equations 2 and 3 allow calculating the equilibrium concentrations of monomers and homomers for the fraction of protein copies that can only homomerize:

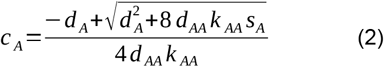

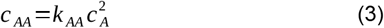

where *c*_*A*_ is the concentration of monomers, *c*_*AA*_ is the concentration of homodimers, *d*_*A*_ and *d*_*AA*_ are the decay rates of monomers and homodimers, *s*_*A*_ is the rate of synthesized copies of protein A that can only homomerize, and *k*_*AA*_ is the association constant for the homodimer. The same equations are used for the equilibrium concentrations for monomers and homodimers of protein B.

Similarly, equations 4 - 8 were used to calculate the equilibrium concentrations for the fraction of paralogs that were allowed to mix:

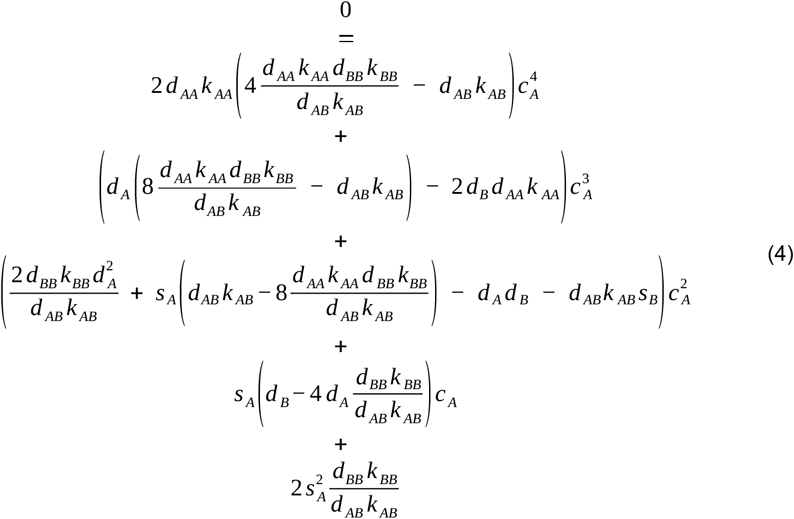

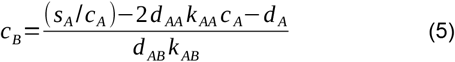

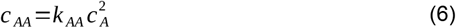

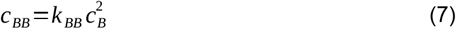

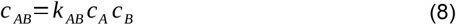

Decay rates were set to 1.3 h^−1^ based on estimates for the half-life of proteins and cell division rates in yeast. The fitness function used 60 h^−1^ as the optimum, allowing us to simulate the case in which it is overshot by the total activity after duplication (76.4 h^−1^). Simulations continued by sampling from the distributions of mutational effects for each of the selected structures until 200 mutations were fixed, with 50 replicates of each simulation. Each sampled mutation affects the folding free energy and the binding energy of the homo- and heterodimers, which modifies their concentrations at equilibrium.

### Annotating barriers to paralog heteromerization

#### Co-/Post-translational assembly data

We used the results from our recently developed framework that integrated structure-based predictions with experimental characterization to gain an overview of co-translational assembly across the *Homo sapiens, Saccharomyces cerevisiae*, and *Escherichia coli* proteome ^111^. In all three organisms, we found that co-translationally assembling complex subunits represent ∼20% of the proteome whereas ∼50% was annotated as post, and the remaining ∼30% could not be annotated due to ambiguous or absent data. These genome-scale annotations were mapped to the 1998 protein families. A protein family was annotated as co- or post-translational if all the mapped family members were either co- or post-translational. This analysis enabled us to annotate 591 co-translational and 792 post-translational families (*Data S3*).

#### Protein subcellular localization data

Subcellular localization data for 13106 human proteins were obtained from the Human Protein Atlas ^105^ and were subsequently filtered to remove uncertain annotations. The final dataset included high-confidence subcellular localization data for 12018 human proteins. Each protein was identified by Ensembl IDs and was mapped onto the human reference proteome according to UniProt mapping (*Data S3*).

#### Gene expression data

We obtained the RNAseq-based presence/absence data of 19619 human transcripts in 257 tissues and 1056 cell types from the Human Protein Atlas ^105^. We also collected the corresponding processed mRNA abundance data (Transcripts per Million, or *TPM* values). The mRNAs were identified by Ensembl IDs and were mapped onto the human reference proteome according to UniProt mapping. In cases where multiple transcripts matched one UniProt ID, only the one with the highest mean abundance was kept (*Data S3*).

#### Interface evolution of paralogous pairs

A three-step approach was undertaken to examine the interface evolution of paralogous pairs.

##### (i) AlphaFold2 models for homomers and heteromers

We compiled a dataset of AlphaFold2-modeled structures of homo- and heteromeric human paralogs. A total of 3760 homomeric models were re- used from our previous study ^71^. Additionally, we generated models of 955 heteromeric pairs using AlphaFold Multimer v2.3 ^106,107^, using a custom ColabFold implementation ^164^. First, for each of the 7377 homomer-derived human paralogous genes, we used MMseqs2 ^165^ to identify their orthologs in 3788 UniProt-annotated eukaryotic proteomes (1^st^ March, 2023). Those with ≥ 80% query sequence coverage (and ≥ 40% target sequence coverage) and 30% ≤ identity ≤ 80% were considered as high-quality hits. The sequence identity upper bound is expected to increase the number of effective sequences from which AlphaFold2 would extract the evolutionary information. From each proteome, we only considered the top hit. The hits for each paralogous pair were then compared, and only those with matching taxon were kept. The resulting hits were further filtered so that no two hits had ≥ 85% sequence identity with each other. The final sets of hits were considered high-quality orthologs. These sequences were then aligned with the Muscle v5.1 using the super5 algorithm ^166^. These alignments were then fed to AlphaFold2, using the monomeric structures from AlphaFold database ^151^ as structural templates. Five heteromeric structures were generated for each of the 2077 paralog pairs, for a total of 10385 models. These models were generated for full-length protein sequences. As a result, these models often contain long flexible, disordered regions masking an otherwise structurally conserved core. These disordered regions were removed to extract these conserved core regions for each complex, using a computational pipeline adapted from Schweke et al. ^71^. This pipeline also computes the probability score for each inferred complex being a physiological dimer. For heteromer, the model with the highest dimer probability was selected. Finally, a total of 878 models with ≥ 0.5 dimer probabilities and core structures of at least 60 residues for both subunits were considered for further analysis, which included 181 *Heteromeric* and 697 *Mixed* pairs (*Data S4*).

##### (ii) Benchmarking the AlphaFold2 models

We used two benchmark tests to assess the accuracy of our heteromer models. First, on May 8th, 2023 we collected experimental structures of 90 heterodimers submitted to PDB after September 30, 2021, the AlphaFold Multimer v2.3 training date. MMseqs2 ^165^ was then used to identify any potential sequence homologs (≥30% identity, ≥60% query coverage, ≥60 aa sequence length) of these structures that were present in the AlphaFold2 training set. We identified a total of 51 benchmark heterodimeric structures with no such potential homologs. These structures were not necessarily heteromeric paralogs. For each benchmark heterodimer, a model structure with dimer probability ≥0.5 was generated following the pipeline described above, and the two structures were subsequently aligned using the MM-align structural alignment tool (version 20191021) ^167^. The reliability of modeled structures was assessed by their RMSD scores of alignment and DockQ scores ^168^. Out of the initial 51, a total of 45 (90%) models accurately recapitulated the heteromeric interaction interface (DockQ ≥ 0.23), with 14 (28%) being high-quality predictions (DockQ ≥ 0.8) (*Figure S5*).

For a second benchmark, we compared our models to those produced by the AlphaFold3 server ^169^. A total of 40 heteromeric paralogous pair models were selected for this analysis, their dimer probabilities spanning the entire range of zero to one. For each, an AlphaFold3 model was generated and then were pairwise compared as described above. Our AlphaFold2 models with dimer probability ≥ 0.5 had low RMSD values and high DockQ scores (*Figure S5*) when compared to their AlphaFold3 counterparts. Thus, our filtered models are comparable to those obtained with AlphaFold3.

These benchmark analyses confirm that our workflow generates and identifies reliable models for heteromeric protein-protein interactions.

##### (iii) Interface analysis. Dimeric interfaces of our high-quality models were annotated following Levy ^170^. Briefly, FreeSASA v2.0.3 ^171^ was used to compute the solvent-accessible surface area (*ASA*) of each amino acid residue in the context of a Gly-X-Gly peptide, in the monomeric (*ASA*_*m*_) and also in the oligomeric state (*ASA*_*o*_). Indeed, *ASA*_*o*_ < *ASA*_*m*_ highlights that the residue is buried at the interface. To examine whether a particular interface residue site is conserved in both paralogs or not, we aligned their sequences using NEEDLE v6.6 ^172^. Comparing the interface residue annotations with the sequence conservation allowed us to measure the variations of the interface. An interface residue site was annotated as ‘conserved’ if it was found at the interface of both paralogs or as ‘novel’ if found at the interface of only one paralog. We used the sequence alignment to identify insertions and deletions (InDels) as well

### Sequence divergence of paralogous pairs

Amino acid sequences of human paralogous pairs were aligned using Clustal Omega ^173^ and Muscle ^166^. These pairwise alignments were used to detect the aligned positions, substitutions, and InDels.

### Structural divergence of paralogous pairs

The core structures of high-quality AF2-derived models were trimmed to their core structures using adapted scripts from Schweke et al. ^71^. The core structures of each pair of paralogs (3703 *Homomeric*, 697 *Mixed*, and 181 *Heteromeric* pairs) were then aligned using the MM-align structural alignment tool (version 20191021) ^167^ to calculate the template-modeling score (TM-score), normalized by the smaller of both paralogs, which is a metric of structural similarity. The TM-score ranges from 0 to 1 and values closer to 1 indicate higher structural similarity.

### Functional divergence of paralogous pairs

#### Enzyme Commission number and reaction data

Enzyme Commission numbers of 3758 annotated human enzymes were obtained from Expasy ^116^ and KEGG databases ^95^. These enzymes mapped to 1622 paralogous pairs (1317 proteins, *Data S3*). An enzyme may have one or more EC numbers assigned to it and the latter includes promiscuous enzyme activity or bifunctional enzymes. For simplicity, we selected only one EC number per enzyme: the one that is the closest to its paralogous enzymes.

#### Gene Ontology data

We obtained Gene Ontology (GO) data for 19070 human proteins from GOA database ^174^ of EMBL-EBI, which included data for 2245 paralogous pairs. Since subcellular compartmentalization acts as a barrier against paralog heteromerization, to avoid circular arguments, GO terms related to subcellular locations were excluded, and those related to ‘biological process’ and ‘molecular function’ were considered in the analysis.

#### Evolutionary rate data

We obtained the rates of the rate of substitutions at synonymous (*dS*) and non-synonymous sites (*dN*) for 12348 human protein-coding genes from a previous systematic analysis of phylogenetic and population data ^129^ (*Data S3*). In this study, the authors compiled comprehensive sequence datasets at phylogenetic scale. The various evolutionary rate parameters were measured by employing a Bayesian Mutation-Selection Framework ^175^. The genes were identified by Ensembl IDs and were mapped onto the human reference proteome according to UniProt mapping.

