## Supplementary_figures for "Co-translational assembly promotes functional diversification of paralogous proteins"

**This Supplementary Materials file includes**

- Supplementary Data File legends (S1-S4)
- Supplementary Tables, S1-S3
- Supplementary Figures, S1-S5
- Supplementary References

### Supplementary Data Files

**Data S1.** The molecular interaction data of human proteins (homomeric and heteromeric).

**Data S2.** The list of orthogroups (OGs), human family members, inferred gene tree, and inferred gene duplication events.

**Data S3.** Subcellular localization, tissue/cell-specific expression, EC numbers, GO terms, and evolutionary rate data for human paralog pairs.

**Data S4.** AlphaFold2 models of paralogous protein complexes are available in Zenodo: https://doi.org/10.5281/zenodo.14158203

### Supplementary Tables

**Table S1.** A set of 123 organisms with fully sequenced genomes included in this study. The phylogenetic classification, scientific name, NCBI taxonomic identifier, UniProt proteome ID, and GenBank Accession ID are provided for each organism.

| Domain | Species | TaxID | Proteome ID | Accession |
|---|---|---|---|---|
| Bacteria | *Escherichia coli* | 83333 | UP000000625 | GCA_000005845.2 |
| Bacteria | *Hahella chejuensis* | 349521 | UP000000238 | GCA_000012985.1 |
| Bacteria | *Pasteurella multocida* | 272843 | UP000000809 | GCA_000006825.1 |
| Bacteria | *Yersinia pestis* | 632 | UP000000815 | GCA_000009065.1 |
| Bacteria | *Ralstonia solanacearum* | 267608 | UP000001436 | GCA_000009125.1 |
| Bacteria | *Albidiferax ferrireducens* | 338969 | UP000008332 | GCA_000013605.1 |
| Bacteria | *Bordetella bronchiseptica* | 257310 | UP000001027 | GCA_000195675.1 |
| Bacteria | *Ruegeria pomeroyi* | 246200 | UP000001023 | GCA_000011965.2 |
| Bacteria | *Caulobacter vibrioides* | 565050 | UP000001364 | GCA_000022005.1 |
| Bacteria | *Rhodospirillum rubrum* | 269796 | UP000001929 | GCA_000013085.1 |
| Bacteria | *Desulfovibrio desulfuricans* | 876 | UP000297065 | GCA_004801255.1 |
| Archaea | *Heimdallarchaeota archaeon strain B3-JM-08* | 2012493 | UP000245584 | GCA_003144275.1 |
| Archaea | *Thorarchaeota archaeon strain OWC* | 2053491 | UP000273160 | GCA_003662775.1 |
| Archaea | *Candidatus Lokiarchaeota archaeon Loki b32* | 2012491 | UP000319869 | GCA_005223125.1 |
| Archaea | *Candidatus Heimdallarchaeota archaeon* | 2026747 | UP000634061 | GCA_014730365.1 |
| Archaea | *Candidatus Helarchaeota archaeon* | 2719382 | UP000625964 | GCA_013375455.1 |
| Archaea | *Candidatus Lokiarchaeota archaeon* | 2053489 | UP000760201 | GCA_011365055.1 |
| Eukaryota | *Galdieria sulphuraria* | 130081 | UP000030680 | GCA_000341285.1 |
| Eukaryota | *Streblomastix strix* | 222440 | UP000324800 | GCA_008636045.1 |
| Eukaryota | *Stylonychia lemnae* | 5949 | UP000039865 | GCA_000751175.1 |
| Eukaryota | *Pocillopora damicornis* | 46731 | UP000275408 | GCA_003704095.1 |
| Eukaryota | *Carpediemonas membranifera* | 201153 | UP000717585 | GCA_019828565.1 |
| Eukaryota | *Allomyces macrogynus* | 578462 | UP000054350 | GCA_000151295.1 |
| Eukaryota | *Conidiobolus coronatus* | 796925 | UP000070444 | GCA_001566745.1 |
| Eukaryota | *Helobdella robusta* | 6412 | UP000015101 | GCA_000326865.1 |
| Eukaryota | *Acanthamoeba castellanii* | 1257118 | UP000011083 | GCA_000313135.1 |
| Eukaryota | *Amphimedon queenslandica* | 400682 | UP000007879 | GCA_000090795.1 |

| | | | | |
|---|---|---|---|---|
| Eukaryota | *Phytophthora infestans* | 403677 | UP000006643 | GCA_000142945.1 |
| Eukaryota | *Perkinsus marinus* | 423536 | UP000007800 | GCA_000006405.1 |
| Eukaryota | *Plasmodiophora brassicae* | 37360 | UP000039324 | GCA_001049375.1 |
| Eukaryota | *Reticulomyxa filosa* | 46433 | UP000023152 | GCA_000512085.1 |
| Eukaryota | *Emiliania huxleyi* | 2903 | UP000013827 | GCA_000372725.1 |
| Eukaryota | *Naegleria gruberi* | 5762 | UP000006671 | GCA_000004985.1 |
| Eukaryota | *Caenorhabditis elegans* | 6239 | UP000001940 | GCA_000002985.3 |
| Eukaryota | *Crassostrea gigas* | 29159 | UP000005408 | GCA_902806645.1 |
| Eukaryota | *Octopus bimaculoides* | 37653 | UP000053454 | GCA_001194135.2 |
| Eukaryota | *Volvox carteri* | 3068 | UP000001058 | GCA_000143455.1 |
| Eukaryota | *Chlamydomonas reinhardtii* | 3055 | UP000006906 | GCA_000002595.3 |
| Eukaryota | *Schistosoma japonicum* | 6182 | UP000311919 | GCA_006368765.1 |
| Eukaryota | *Echinococcus multilocularis* | 6211 | UP000017246 | GCA_000469725.3 |
| Eukaryota | *Leishmania major* | 5664 | UP000000542 | GCA_000002725.2 |
| Eukaryota | *Trypanosoma brucei* | 185431 | UP000008524 | GCA_000002445.1 |
| Eukaryota | *Phaeodactylum tricornutum* | 556484 | UP000000759 | GCA_000150955.2 |
| Eukaryota | *Thalassiosira pseudonana* | 35128 | UP000001449 | GCA_000149405.2 |
| Eukaryota | *Plasmodium falciparum* | 36329 | UP000001450 | GCA_000002765.3 |
| Eukaryota | *Toxoplasma gondii* | 432359 | UP000002226 | GCA_000150015.2 |
| Eukaryota | *Ustilago maydis* | 237631 | UP000000561 | GCA_000328475.2 |
| Eukaryota | *Sporisorium reilianum* | 999809 | UP000008867 | GCA_000230245.1 |
| Eukaryota | *Jaapia argillacea* | 933084 | UP000027265 | GCA_000697665.1 |
| Eukaryota | *Fomitopsis pinicola* | 743788 | UP000015241 | GCA_000344655.2 |
| Eukaryota | *Calocera cornea* | 1353952 | UP000076842 | GCA_001632435.1 |
| Eukaryota | *Imshaugia aleurites* | 172621 | UP000664534 | GCA_905337355.1 |
| Eukaryota | *Botryotinia fuckeliana* | 332648 | UP000001798 | GCA_000143535.4 |
| Eukaryota | *Magnaporthe grisea* | 148305 | UP000515153 | GCF_004355905.1 |
| Eukaryota | *Colletotrichum fructicola* | 1213859 | UP000011096 | GCA_000319635.2 |
| Eukaryota | *Alternaria alternata* | 5599 | UP000077248 | GCA_001642055.1 |
| Eukaryota | *Zymoseptoria tritici* | 1276538 | UP000215127 | GCA_900091695.1 |
| Eukaryota | *Exophiala dermatitidis* | 858893 | UP000007304 | GCA_000230625.1 |
| Eukaryota | *Aspergillus clavatus* | 344612 | UP000006701 | GCA_000002715.1 |
| Eukaryota | *Saccharomyces cerevisiae* | 559292 | UP000002311 | GCA_000146045.2 |
| Eukaryota | *Debaryomyces hansenii* | 284592 | UP000000599 | GCA_000006445.2 |

| | | | | |
|---|---|---|---|---|
| Eukaryota | *Arabidopsis thaliana* | 3702 | UP000006548 | GCA_000001735.1 |
| Eukaryota | *Brachypodium distachyon* | 15368 | UP000008810 | GCA_000005505.4 |
| Eukaryota | *Amborella trichopoda* | 13333 | UP000017836 | GCA_000471905.1 |
| Eukaryota | *Macleaya cordata* | 56857 | UP000195402 | GCA_002174775.1 |
| Eukaryota | *Selaginella moellendorffii* | 88036 | UP000001514 | GCA_000143415.2 |
| Eukaryota | *Marchantia polymorpha* | 1480154 | UP000077202 | GCA_001641455.1 |
| Eukaryota | *Dermatophagoides pteronyssinus* | 6956 | UP000515146 | GCF_001901225.1 |
| Eukaryota | *Varroa destructor* | 109461 | UP000594260 | GCA_002443255.1 |
| Eukaryota | *Drosophila melanogaster* | 7227 | UP000000803 | GCA_000001215.4 |
| Eukaryota | *Pediculus humanus* | 121224 | UP000009046 | GCA_000006295.1 |
| Eukaryota | *Aphis craccivora* | 307492 | UP000478052 | GCA_009835225.1 |
| Eukaryota | *Apis mellifera* | 7460 | UP000005203 | GCF_003254395.2 |
| Eukaryota | *Strigamia maritima* | 126957 | UP000014500 | GCA_000239455.1 |
| Eukaryota | *Darwinula stevensoni* | 69355 | UP000677054 | GCA_905338385.1 |
| Eukaryota | *Homarus americanus* | 6706 | UP000747542 | GCA_018991925.1 |
| Eukaryota | *Erpetoichthys calabaricus* | 27687 | UP000694620 | GCA_900747795.2 |
| Eukaryota | *Callorhinchus milii* | 7868 | UP000314986 | GCA_000165045.2 |
| Eukaryota | *Ciona intestinalis* | 7719 | UP000008144 | GCA_000224145.1 |
| Eukaryota | *Geotrypetes seraphini* | 260995 | UP000515159 | GCF_902459505.1 |
| Eukaryota | *Xenopus laevis* | 8355 | UP000186698 | GCF_017654675.1 |
| Eukaryota | *Branchiostoma belcheri* | 7741 | UP000515135 | GCF_001625305.1 |
| Eukaryota | *Sphenodon punctatus* | 8508 | UP000694392 | GCA_003113815.1 |
| Eukaryota | *Podarcis muralis* | 64176 | UP000472272 | GCA_004329235.1 |
| Eukaryota | *Anolis carolinensis* | 28377 | UP000001646 | GCA_000090745.2 |
| Eukaryota | *Neogobius melanostomus* | 47308 | UP000694523 | GCA_007210695.1 |
| Eukaryota | *Chanos chanos* | 29144 | UP000504632 | GCF_902362185.1 |
| Eukaryota | *Gadus morhua* | 8049 | UP000694546 | GCA_902167405.1 |
| Eukaryota | *Electrophorus electricus* | 8005 | UP000314983 | GCA_003665695.2 |
| Eukaryota | *Sphaeramia orbicularis* | 375764 | UP000472271 | GCA_902148855.1 |
| Eukaryota | *Salmo salar* | 8030 | UP000087266 | GCF_000233375.1 |
| Eukaryota | *Lepisosteus oculatus* | 7918 | UP000018468 | GCA_000242695.1 |
| Eukaryota | *Hippocampus comes* | 109280 | UP000264820 | GCA_001891065.1 |
| Eukaryota | *Oryzias latipes* | 8090 | UP000001038 | GCA_002234675.1 |
| Eukaryota | *Pteropus vampyrus* | 132908 | UP000515202 | GCF_000151845.1 |

| | | | | |
|---|---|---|---|---|
| Eukaryota | *Phyllostomus discolor* | 89673 | UP000504628 | GCF_004126475.2 |
| Eukaryota | *Sarcophilus harrisii* | 9305 | UP000007648 | GCA_902635505.1 |
| Eukaryota | *Monodelphis domestica* | 13616 | UP000002280 | GCA_000002295.1 |
| Eukaryota | *Phascolarctos cinereus* | 38626 | UP000515140 | GCF_002099425.1 |
| Eukaryota | *Erinaceus europaeus* | 9365 | UP000079721 | GCF_000296755.1 |
| Eukaryota | *Oryctolagus cuniculus* | 9986 | UP000001811 | GCA_000003625.1 |
| Eukaryota | *Ornithorhynchus anatinus* | 9258 | UP000002279 | GCA_004115215.2 |
| Eukaryota | *Equus caballus* | 9796 | UP000002281 | GCA_002863925.1 |
| Eukaryota | *Loxodonta africana* | 9785 | UP000007646 | GCA_000001905.1 |
| Eukaryota | *Mus musculus* | 10090 | UP000000589 | GCA_000001635.9 |
| Eukaryota | *Ictidomys tridecemlineatus* | 43179 | UP000005215 | GCA_000236235.1 |
| Eukaryota | *Marmota marmota* | 9994 | UP000694407 | GCA_001458135.1 |
| Eukaryota | *Cavia porcellus* | 10141 | UP000005447 | GCA_000151735.1 |
| Eukaryota | *Trichechus manatus* | 127582 | UP000248480 | GCF_000243295.1 |
| Eukaryota | *Bos taurus* | 9913 | UP000009136 | GCA_002263795.2 |
| Eukaryota | *Sus scrofa* | 9823 | UP000008227 | GCA_000003025.6 |
| Eukaryota | *Tursiops truncatus* | 9739 | UP000245320 | GCF_011762595.1 |
| Eukaryota | *Vicugna pacos* | 30538 | UP000504605 | GCF_000164845.4 |
| Eukaryota | *Canis lupus* | 9615 | UP000002254 | GCA_000002285.4 |
| Eukaryota | *Felis catus* | 9685 | UP000011712 | GCA_000181335.4 |
| Eukaryota | *Panthera tigris* | 74533 | UP000675900 | GCA_000464555.1 |
| Eukaryota | *Ursus maritimus* | 29073 | UP000261680 | GCF_017311325.1 |
| Eukaryota | *Homo sapiens* | 9606 | UP000005640 | GCA_000001405.29 |
| Eukaryota | *Gorilla gorilla* | 9595 | UP000001519 | GCA_000151905.3 |
| Eukaryota | *Macaca mulatta* | 9544 | UP000006718 | GCA_003339765.3 |
| Eukaryota | *Pygocentrus nattereri* | 42514 | UP000261440 | GCA_001682695.1 |
| Eukaryota | *Ictalurus punctatus* | 7998 | UP000221080 | GCF_001660625.3 |
| Eukaryota | *Danio rerio* | 7955 | UP000000437 | GCF_000002035.6 |

**Table S2.** The increase in %heteromers in the *final equilibrium*, as compared to that in the *initial equilibrium*, for the 37 structures, for various separation efficiency coefficients. Two-sample t-test statistics and the corresponding *p*-values are provided.

| Separation efficiency | %Heteromers | | | t-statistics | *p*-value |
|---|---|---|---|---|---|
| | initial equilibrium | final equilibrium | Δ (final, initial) | | |
| 0.0 | 50.0 | 69.0 | 19.0 | 7.5 | 3.30E-09 |
| 0.1 | 45.0 | 60.0 | 15.0 | 7.8 | 1.40E-09 |
| 0.2 | 40.0 | 55.0 | 15.0 | 8.2 | 5.00E-10 |
| 0.3 | 35.0 | 47.0 | 12.0 | 7.4 | 4.50E-09 |
| 0.4 | 30.0 | 41.0 | 11.0 | 8.7 | 9.90E-11 |
| 0.5 | 25.0 | 34.1 | 9.1 | 7.7 | 2.00E-09 |
| 0.6 | 20.0 | 27.1 | 7.1 | 7.0 | 1.50E-08 |
| 0.7 | 15.0 | 20.3 | 5.3 | 7.3 | 7.20E-09 |
| 0.8 | 10.0 | 13.6 | 3.6 | 7.5 | 3.70E-09 |
| 0.9 | 5.0 | 6.6 | 1.6 | 5.7 | 8.00E-07 |
| 1.0 | 0.0 | 0.0 | 0.0 | -Inf | 1.00E+00 |

**Table S3.** The proportions of heteromers observed after removing the barrier *versus* those obtained for simulations with no separation are compared by two-sample t-tests. Each row corresponds to the PDB identifier of the simulated complex, each column represents a certain separation efficiency threshold. The numbers denote the *p*-value of the corresponding t-test.

| PDB | q = 0 | q = 0.1 | q = 0.2 | q = 0.3 | q = 0.4 | q = 0.5 | q = 0.6 | q = 0.7 | q = 0.8 | q = 0.9 | q = 1.0 |
|---|---|---|---|---|---|---|---|---|---|---|---|
| 1a4b | 0.95 | 0.47 | 0.03 | 0.19 | 0.13 | 0.01 | 0.09 | 0.16 | 0.01 | 0.09 | 0.38 |
| 1ai2 | 0.97 | 0.60 | 0.04 | 0.04 | 0.13 | 0.51 | 0.12 | 0.45 | 0.32 | 0.15 | 0.53 |
| 1asb | 0.90 | 0.89 | 0.80 | 0.57 | 0.82 | 0.78 | 0.95 | 0.67 | 0.85 | 0.25 | 0.92 |
| 1b57 | 0.66 | 0.17 | 0.70 | 0.54 | 0.39 | 0.91 | 0.14 | 0.13 | 0.45 | 0.18 | 0.93 |
| 1bbh | 0.90 | 0.60 | 0.18 | 0.90 | 0.83 | 0.21 | 0.41 | 0.87 | 0.43 | 0.02 | 0.98 |
| 1btm | 0.84 | 0.18 | 0.01 | 0.32 | 0.01 | 0.01 | 0.01 | 0.04 | 0.17 | 0.68 | 0.07 |
| 1c1f | 0.82 | 0.02 | 0.37 | 0.03 | 0.00 | 0.03 | 0.14 | 0.12 | 0.00 | 0.10 | 0.17 |
| 1eye | 0.84 | 0.45 | 0.43 | 0.56 | 0.69 | 0.98 | 0.45 | 0.19 | 0.23 | 0.32 | 0.66 |
| 1g99 | 0.96 | 0.79 | 0.71 | 0.05 | 0.79 | 0.91 | 0.30 | 0.72 | 0.69 | 0.71 | 0.05 |
| 1gpr | 1.00 | 0.47 | 0.79 | 0.60 | 0.36 | 0.36 | 0.72 | 0.80 | 0.75 | 0.88 | 0.36 |
| 1h0c | 0.68 | 0.82 | 0.92 | 0.68 | 0.09 | 0.89 | 0.93 | 0.99 | 0.63 | 0.93 | 0.52 |
| 1i49 | 0.98 | 0.84 | 0.82 | 0.35 | 0.84 | 0.97 | 0.81 | 0.14 | 0.75 | 0.60 | 0.48 |
| 1m38 | 0.99 | 0.11 | 0.16 | 0.49 | 0.09 | 0.07 | 0.73 | 0.29 | 0.28 | 0.94 | 0.88 |
| 1m7p | 0.93 | 0.08 | 0.38 | 0.39 | 0.52 | 0.13 | 0.12 | 0.22 | 0.20 | 0.87 | 0.51 |
| 1mk4 | 0.97 | 0.85 | 0.43 | 0.18 | 0.25 | 0.82 | 0.38 | 0.94 | 0.14 | 0.78 | 0.94 |
| 1nmz | 1.00 | 0.86 | 0.98 | 0.36 | 0.92 | 0.66 | 0.98 | 0.88 | 0.31 | 0.54 | 0.50 |
| 1p6o | 0.32 | 0.18 | 0.64 | 0.84 | 0.97 | 0.39 | 0.64 | 0.16 | 0.11 | 0.07 | 0.09 |
| 1pzw | 0.90 | 0.84 | 0.59 | 0.12 | 0.73 | 0.71 | 0.63 | 0.38 | 0.65 | 0.05 | 0.38 |
| 1uuf | 0.92 | 0.24 | 0.01 | 0.14 | 0.73 | 0.09 | 0.08 | 0.47 | 0.25 | 0.08 | 0.26 |
| 2b18 | 1.00 | 0.24 | 0.03 | 0.37 | 0.44 | 0.05 | 0.21 | 0.48 | 0.07 | 0.10 | 0.21 |
| 2pih | 0.93 | 0.16 | 0.79 | 0.55 | 0.87 | 0.43 | 0.94 | 0.57 | 0.76 | 0.56 | 0.34 |
| 2raj | 0.89 | 0.85 | 0.91 | 0.59 | 0.24 | 0.22 | 0.12 | 0.50 | 0.21 | 0.10 | 0.44 |
| 2vtk | 0.98 | 0.65 | 0.17 | 0.19 | 0.45 | 0.17 | 0.17 | 0.04 | 0.05 | 0.12 | 0.46 |
| 3euo | 0.91 | 0.08 | 0.38 | 0.95 | 0.30 | 0.78 | 0.61 | 0.56 | 0.05 | 0.49 | 0.75 |
| 3f4d | 0.86 | 0.11 | 0.59 | 0.81 | 0.72 | 0.63 | 0.99 | 0.81 | 0.46 | 0.91 | 0.71 |
| 3gi4 | 0.21 | 0.56 | 0.01 | 0.91 | 0.18 | 0.03 | 0.59 | 0.01 | 0.42 | 0.71 | 0.37 |
| 3h65 | 0.89 | 0.69 | 0.93 | 0.18 | 0.37 | 0.10 | 0.63 | 0.19 | 0.29 | 0.78 | 0.73 |
| 3ib7 | 0.80 | 0.02 | 0.42 | 0.91 | 0.96 | 0.02 | 0.91 | 0.45 | 0.40 | 0.54 | 0.45 |
| 3naq | 0.90 | 0.10 | 0.88 | 0.24 | 0.93 | 0.96 | 0.49 | 0.53 | 0.89 | 0.77 | 0.80 |
| 3u2g | 1.00 | 0.04 | 0.02 | 0.52 | 0.93 | 0.53 | 0.63 | 0.09 | 0.16 | 0.11 | 0.86 |

| | | | | | | | | | | | |
|---|---|---|---|---|---|---|---|---|---|---|---|
| 3zof | 0.54 | 0.65 | 0.84 | 0.04 | 0.87 | 0.54 | 0.46 | 0.70 | 0.14 | 0.26 | 0.32 |
| 4cwd | 0.79 | 0.11 | 0.75 | 0.92 | 0.56 | 0.05 | 0.31 | 0.53 | 0.72 | 0.90 | 0.99 |
| 4fgw | 0.90 | 0.83 | 0.81 | 0.32 | 0.55 | 0.18 | 0.78 | 0.67 | 0.97 | 0.30 | 0.73 |
| 4o48 | 0.92 | 0.51 | 0.95 | 0.54 | 0.84 | 0.86 | 0.66 | 0.23 | 0.75 | 0.77 | 0.92 |
| 4wpe | 0.97 | 0.08 | 0.48 | 0.20 | 0.15 | 0.03 | 0.87 | 0.64 | 0.15 | 0.10 | 0.16 |
| 5bj4 | 0.49 | 0.94 | 0.56 | 0.75 | 0.97 | 0.75 | 0.84 | 0.68 | 0.94 | 0.73 | 0.75 |
| 5hbi | 0.91 | 0.69 | 0.71 | 0.61 | 0.23 | 0.08 | 0.61 | 0.85 | 0.16 | 0.66 | 0.67 |

#### Supplementary Figures

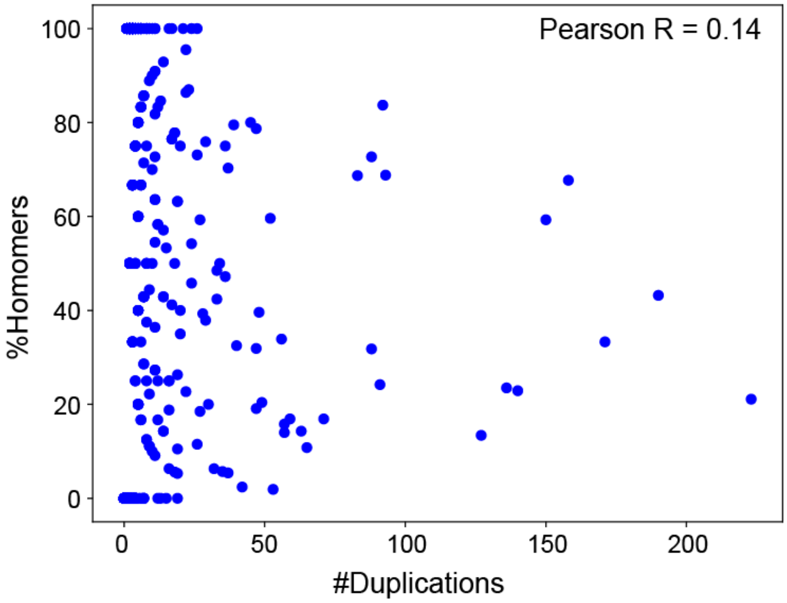


**Figure S1. The proportion of homomeric/heteromeric fates does not strongly correlate with the frequency of duplications.** The X-axis represents the frequency of gene duplications at each ancestral node of the eukaryotic Tree of Life; the Y-axis represents the percent of those duplications that gave rise to *Homomeric* paralogous pairs.

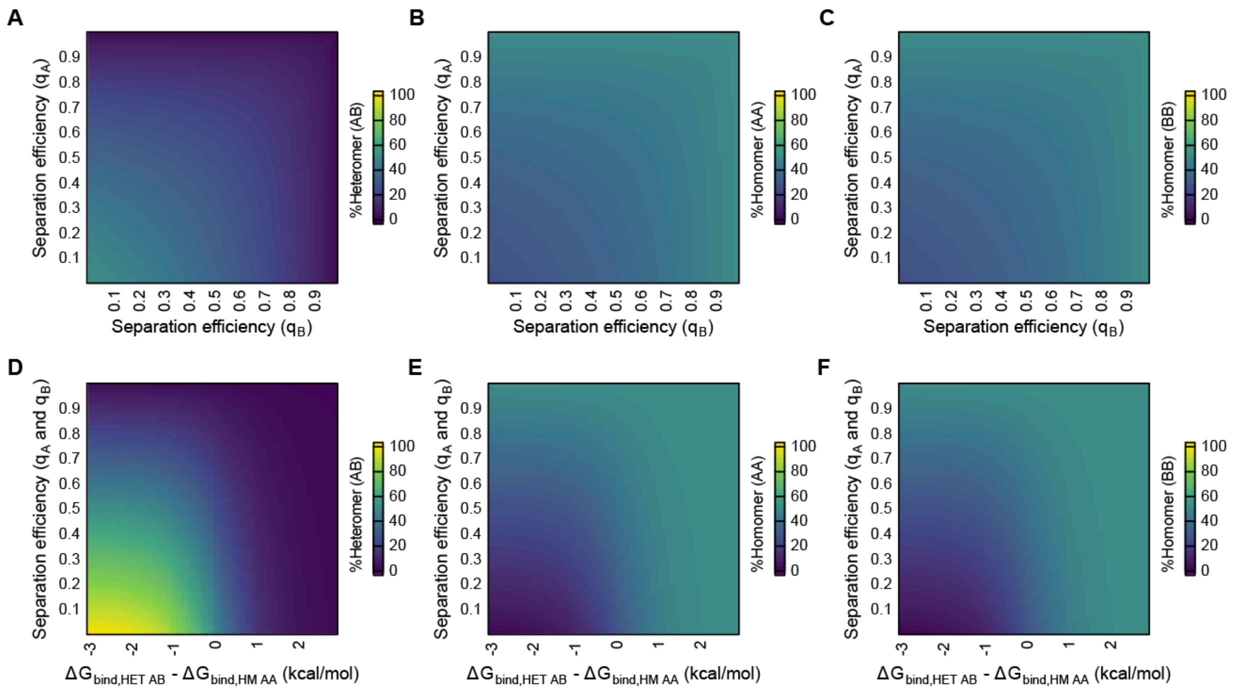


**Figure S2. Introducing separation efficiencies constrains the proportion of heterodimers in the simulations.** (**A-C**) The percentages of the heterodimer (panel **A**) and that of each of the two homodimers (panels **B**, and **C**) as a function of the separation efficiencies assigned to each of the two protein subunits. For two paralogs A and B, homomers and heteromers are denoted as AA, BB, and AB. In all three panels, binding affinities are identical for the three dimers. As separation efficiency increases, heteromers are depleted and the proportion of homomers increases. (**D-F**) Percentages of heterodimers (panel **D**) and that of each of the two homodimers (panels **E**, and **F**) as a function of both separation efficiency and binding affinity. HM: Homomer, HET: Heteromer. In all three panels, the binding affinities of the two homodimers are kept constant, while that of the heterodimer was varied. The x-axis represents the Δ binding affinity between the heteromer and the two homomers. The y-axis represents the separation efficiency which is identical for both paralogs.

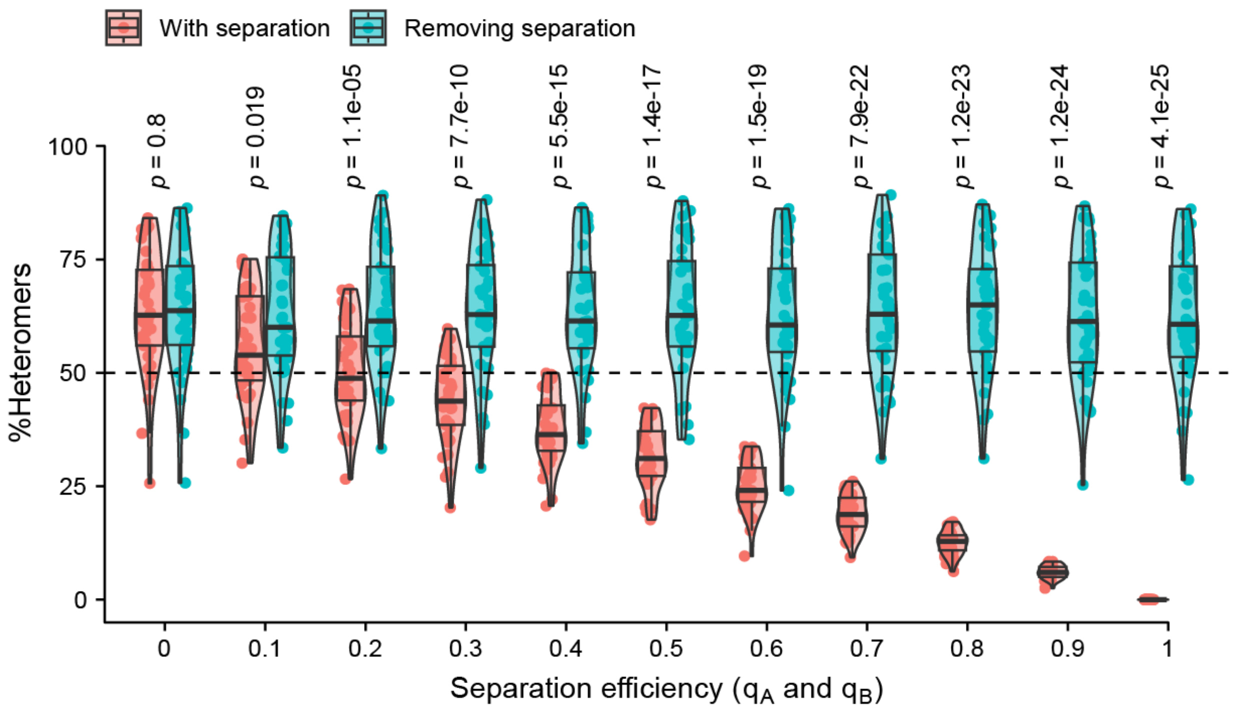


**Figure S3. Paralogs evolving with complete separation may not lose the capacity to self-interact when the barrier is removed, given that the interface remains constant.** The red violin plots represent the %heterodimers in the *final equilibrium* for different separation efficiencies. Each dot represents the average %heterodimers obtained for one complex over 50 simulation replicates. A total of 37 high-quality dimeric complexes were analyzed. The blue violin plots represent the same but were computed using the folding free energies of each paralog and the binding affinities of their complexes obtained in the *final equilibrium*, assuming no separation. The *p*-values were derived from two-sample t-tests.

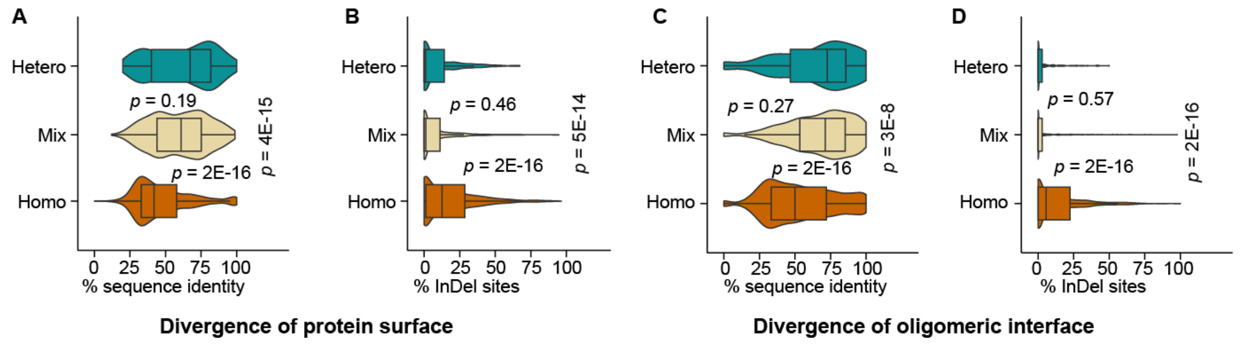


**Figure S4. Sequence divergence of paralogs's solvent-accessible surface and oligomeric interface.** Two paralogs may diverge by accumulating substitutions or InDels. Pairwise sequence alignments were generated for each paralogous pair and the aligned positions and InDel positions were identified. (**A-B**) Sequence divergence of paralogs' solvent-accessible surface. (**A**) Violin plots represent the %sequence identity (representing substitution accumulation) of *Homomeric* (orange), *Heteromeric* (green), and *Mixed* (yellow) paralogous pairs. The *p*-values were derived from pairwise t-tests. (**B**) Same as panel-**A**, for %alignment positions that are InDels. (**C-D**) Same as **A-B**, for sequence divergence of paralogs' oligomeric interface.

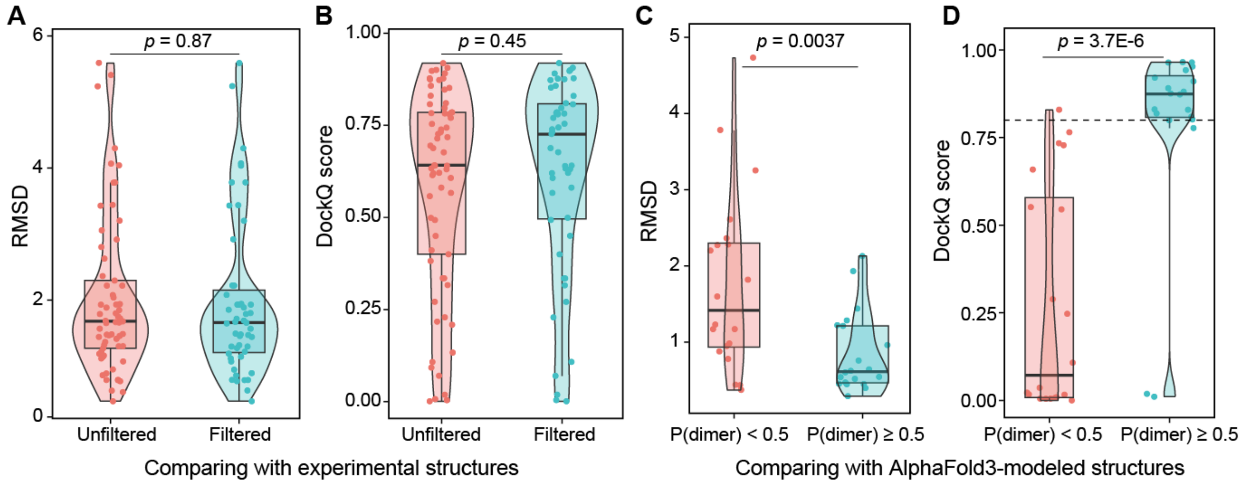


**Figure S5. Quality assessment statistics for the AlphaFold2 benchmark set.** (**A-B**) Comparing the AlphaFold2-modeled structures with experimental structures available in Protein Data Bank (PDB). The root-mean-square distance (RMSD, panel **A**) of the structural alignment of the model and the experimental structures, and the DockQ score of the model structure (panel **B**) were considered as quality measures. The *p*-values were derived from pairwise t-tests. The unfiltered benchmark set (red) includes 65 pairs of interacting proteins submitted to the PDB after the AF2 training date (September 30, 2021). The filtered set (blue) contains the 51 high-quality AF2 models (dimer probability ≥ 0.5) evaluated with adapted scripts from Schweke et al. [1]. (**C-D**) Same as panels **A-B**, for comparing the AlphaFold2-modeled structures with AlphaFold3-modeled structures of the same heterodimeric complexes. AF2 models were classified according to the dimer probabilities (x-axis). The horizontal dashed line in panel B indicates a DockQ score of 0.8, which is typically used to indicate high-quality models [2].
